# Loss of function of chromatin remodeler OsCLSY4 leads to RdDM-mediated mis-expression of endosperm-specific genes affecting grain qualities

**DOI:** 10.1101/2025.07.07.663415

**Authors:** Avik Kumar Pal, Saumyapriya Rana, Riju Dey, P. V. Shivaprasad

**Affiliations:** National Centre for Biological Sciences, Tata Institute of Fundamental Research, GKVK Campus, Bangalore 560 065, India

**Keywords:** Endosperm, RNA silencing, Epigenetics, DNA methylation, small RNAs, Chromatin remodeler, CLSY genes

## Abstract

RNA-directed DNA methylation (RdDM) sequence-specifically targets transposable elements (TEs) and repeats in plants, often in a tissue-specific manner. In triploid endosperm tissue, RdDM also acts as a parental dosage regulator, mediating spatio-temporal expression of genes required for its development. It is unclear how RdDM is initiated and established in endosperm. Rice endosperm-specific imprinted chromatin remodeler OsCLSY3 recruits RNA polymerase IV to specific genomic sites for silencing and optimal gene expression. Here we show that, in addition to OsCLSY3, ubiquitously expressed OsCLSY4 is also crucial for proper reproductive growth and endosperm development. Loss of function of OsCLSY4 led to reproductive and nutrient-filling defects in endosperm. Using genetic and molecular analysis, we show that OsCLSY3 and OsCLSY4 play both overlapping and unique silencing roles in rice endosperm by targeting specific and common TEs, repeats and genic regions. These results indicate the importance of optimal expression of two OsCLSYs in regulating endosperm-specific gene expression, genomic imprinting and suppression of specific TEs. Results presented here provide new insights into the functions of rice CLSYs as upstream RdDM regulators in rice endosperm development, and we propose that functions of their homologs might be conserved across monocots.

## Introduction

In addition to the embryo, angiosperm seed contains a seed coat that protects the embryo from damages, unfavorable conditions and a nutrition-tissue called endosperm that provides nutrients during development (Baroux et al. 2002; Pires 2014). Unlike gymnosperms, majority of the angiosperm seeds contain a triploid endosperm which is also a product of fertilization similar to embryo (Baroux et al. 2002; Gehring 2013; Wang and Köhler 2017). Most of the flowering plants undergo double fertilization events, where one of the two sperm cells fertilizes the egg cell to generate diploid embryo and the other sperm cell fertilizes pre-fused 2n central cell to produce triploid endosperm. This kind of fertilization specific to flowering plants is called double fertilization (Berger et al. 2008). Incorporation of paternal genome in the nutrient tissue endosperm might have helped better allocation of nutrients to the growing embryo, and might have contributed to the rapid evolution of angiosperms often considered as an abominable mystery (De Jong et al. 2005; Lafon-Placette and Köhler 2016).

In cereal seeds, starch-filled endosperm is the major food source for humans and other animals (Li and Berger 2012; Gehring 2013). Its parental-genomic dose, ploidy and chromatin organization are unique when compared to other plant tissues. The molecular mechanism of endosperm development is not yet fully understood (Li and Berger 2012). Besides acting as a source of nutrients for the embryo, endosperm also senses environmental changes, abiotic and biotic factors to regulate seed germination (Chahtane et al. 2018; Iwasaki et al. 2019, 2022; De Giorgi et al. 2021). Multiple studies in model plant *Arabidopsis* indicated that hormonal, genetic and epigenetic pathways regulate gene expression during endosperm development (Baroux et al. 2002; Gehring 2013).

Among these mechanisms, epigenetic changes such as DNA methylation, demethylation and histone modifications are important because they also regulate maternal and paternal genome dose in endosperm (Köhler et al. 2003; Jullien et al. 2006; Köhler and Weinhofer-Molisch 2010; Holec and Berger 2012; Gehring 2013; Satyaki and Gehring 2017). Several upstream and downstream players in these pathways are well-known especially in *Arabidopsis* (Grossniklaus et al. 1998; Xiao et al. 2003; Jullien et al. 2006; Erdmann et al. 2017; Wang et al. 2020). These include players involved in DNA methylation, small RNA (sRNA) biogenesis and targeting of specific genomic sites for epigenetic modifications. Cytosine methylation which is observed in CG, CHG and CHH contexts (where H corresponds to A, T, or C) in plants requires various methyltransferases such as METHYLTRANSFERASE1 (MET1) and CHROMOMETHYLASE3 (CMT3), that regulate CG and CHG methylation, respectively. The CHH methylation is regulated by a *de novo* DNA methyltransferase named DOMAINS REARRANGED METHYLTRANSFERASE2 (DRM2), which is guided by sRNAs. In many selective loci, CHH methylation is also regulated by CHROMOMETHYLASE2 (CMT2) (Stroud et al. 2014). An RNA-mediated regulatory loop is required for initiating and maintaining DNA methylation. A plant-specific DNA-dependent RNA polymerase IV (Pol IV) generates short single-stranded (ss) RNA transcripts predominantly from TEs and repeats. These ssRNAs get converted into double-stranded (ds) forms by RNA DEPENDENT RNA POLYMERASE2 (RDR2) and further processed into 24 nt sRNA duplexes by DICER-LIKE3 (DCL3). The 24 nt sRNAs are preferentially loaded into ARGONAUTE4/6/9 (AGO4/6/9) to guide DRM2 to methylate TEs and repeats by associating with long non-coding transcripts generated by plant-specific DNA-dependent RNA polymerase V (Pol V). Recruitment of Pol IV at specific loci requires a family of putative SNF2 chromatin remodelers named CLASSYs (CLSY), sometimes along with SAWADEE HOMEODOMAIN HOMOLOG1 (SHH1) proteins. The tissue-specific expression pattern of CLSYs regulate tissue-specific DNA methylation in *Arabidopsis* (Smith et al. 2007; Law et al. 2011; Zhang et al. 2013; Zhou et al. 2018, 2022; Martins and Law 2023). Mutations in these genes affect DNA methylation level, plant growth and development, sometimes specifically in the reproductive stages (Walker et al. 2018; Wang et al. 2020; Jha et al. 2021; Long et al. 2021; Chakraborty et al. 2022; Pal et al. 2024).

Most of the mechanistic understanding about CLSY proteins is from *Arabidopsis*. *Arabidopsis* has four CLSY proteins and they are major upstream regulators of the RdDM pathway. The *clsy* quadruple mutant shows global loss of 24 nt sRNAs, similar to *pol iv* mutant (Zhou et al. 2018; Felgines et al. 2024). The sRNA production and DNA methylation were affected in each single CLSY mutants, which suggested they are largely non-overlapping and locus-specific regulators (Zhou et al. 2018, 2022). Although, *clsy1,2* and *clsy3,4* double mutants regulated comparatively more number of sRNA loci than any of the single mutants suggesting that there is also some degree of redundancy within the CLSY family proteins (Zhou et al. 2018, 2022; Xu and Law 2024). Strikingly, in a genetic screen, *Arabidopsi*s CLSY4 was identified as a DNA demethylation factor and strong hypermethylation at several loci was observed in clsy4 (Yang et al. 2018). The study further showed that all single *clsy* mutants displayed many hypermethylated DNA loci along with RdDM-dependent hypomethylated sites, mechanistic understanding of which was not understood (Yang et al. 2018; Pal et al. 2024). Unlike *Arabidopsis*, CLSY mis-expression lines in rice displayed partial or complete sterility due to abnormal reproductive development and seed-filling defects (Xu et al. 2023; Pal et al. 2024).

Endosperm development is quite distinct in dicots compared to monocots. In mature dicot seeds, endosperm tissue gets used up during embryo development, but in monocots, endosperm gets retained and utilized during germination. However, key regulators of endosperm development among monocots are not fully identified yet. While RdDM pathway seems to play an important role in rice reproduction as seen in different mutants based on the phenotypes (Xu et al. 2020, 2023; Chakraborty et al. 2022; Wang et al. 2022; Hari Sundar G et al. 2023; Pal et al. 2024), clear mechanistic understanding has eluded. Rice has three CLSY genes. OsCLSY3 is an endosperm-preferred gene that regulates a set of imprinted genes in endosperm *via* imprinted OsCLSY3-dependent sRNAs (Pal et al. 2024). OsCLSY1 appears to be a crucial gene during anaerobic germination (Castano-Duque et al. 2021). OsCLSY4 was identified as an important RdDM pathway gene in a genetic mutant screening where it was named as *fem2* (Xu et al. 2023). *OsCLSY4* (named here *FEM2*) is a ubiquitously expressed gene important for the regulation of a majority of TEs in vegetative stages. This study further found that in fem2 seedlings, overexpression of *OsCLSY3* (*FEL1*), but not *OsCLSY1* (*FEL2*) partially complemented DNA methylation levels at some specific genomic loci, indicating a partial redundancy in functions between *OsCLSY3* and *OsCLSY4* (Xu et al. 2023).

In this study, we identified redundant and unique RdDM-associated functions of two *OsCLSY* genes that express in the endosperm, a tissue largely exhibiting DNA hypomethylation. We show that in addition to OsCLSY3, OsCLSY4 also redundantly regulated RdDM at specific genomic loci in the endosperm. RNA-seq, sRNA-seq and whole genome DNA methylation analysis indicated that both OsCLSY proteins regulate the same family of TEs. We also identified numerous loci where we observed hypermethylation in the absence of CLSYs. These results indicate that endosperm-preferred OsCLSY3, as well as ubiquitously expressed OsCLSY4 are both crucial for endosperm development through DNA methylation. Significant numbers of seed development, endosperm development, as well as imprinted genes were mis-expressed in their knockdown (kd) lines, providing a molecular basis for the fertility/seed development defects observed in transgenic lines. These findings support the presence of an interplay between redundant and parallel epigenetic pathways mediated by OsCLSY3 and OsCLSY4 proteins in mediating the cellularization timing and development of endosperm tissue in cereals.

## Results

### OsCLSY4 loss of function led to reduced fertility and increased seed defects

In our previous study, we identified maternally expressed OsCLSY3 as an important regulator of the endosperm-specific RdDM pathway in rice. The OsCLSY3-dependent sRNAs regulated expression of multiple seed development associated and imprinted genes *via* DNA methylation. OsCLSY3 expression was also controlled by RdDM pathway in vegetative tissues due to the presence of two MITE TEs in its promoter that are silenced by ubiquitously expressed CLSY homolog OsCLSY4 (Pal et al. 2024). We also observed the expression of many sRNA loci that are independent of OsCLSY3 in the endosperm that was partially attributable to redundancy with other CLSY members (Pal et al. 2024). We observed hyper methylation at several genomic loci, indicating competition with another CLSY or other players involved in RdDM. In endosperm tissue, OsCLSY4 is also expressed (Supplemental Figure S1A, B) while OsCLSY1 is not expressed, indicating a possible redundancy or competition with ubiquitously expressed OsCLSY4 (Pal et al. 2024). We found that both OsCLSY3 and OsCLSY4 possess N-terminal intrinsically-disorder regions, C-terminal SNF2, and helicase domains. They had 31.3% sequence identity and 43.4% similarity between themselves. The N-terminal region of these proteins was comparatively less conserved but C-terminal regions (1030-1440 amino acids) were very well-conserved. These comparisons indicate possibilities of redundant as well as unique functions for these OsCLSYs. We also compared OsCLSY4 protein with CLSY homolog of maize (RMR1) and found that it exhibited around 64.4% sequence identity and 73.4% of similarity. In maize, RMR1 is an important RdDM pathway component and a regulator of paramutation (Hale et al. 2007). In previous studies, knockout (ko) of OsCLSY3 and OsCLSY4 led to partial or complete sterility (Xu et al. 2023; Pal et al. 2024). All these pointed out to a possibility that OsCLSY4 might also be playing an important role in rice endosperm development.

To understand the functional redundancy or unique roles of OsCLSY3 and OsCLSY4 in rice endosperm, and to compare with clsy3-kd data, we used an efficient artificial miRNA (amiR) based kd method to silence OsCLSY4 in *indica* rice line Pusa Basmati-1 (henceforth, PB1) (Narjala et al. 2020). We found up to 70% reduction in *OsCLSY4* transcripts in kd transgenic lines in RT-qPCR analysis (Fig. 1A). As observed in CLSY4/ fem2 mutant, these osclsy4-kd plants had reduced vegetative growth (Supplemental Figure S1C, D). Most importantly, osclsy4-kd lines showed many reproductive defects, such as in panicle length, number of spikelets and grain filling rate per panicle, all of which were drastically reduced (Fig. 1B, C). The RdDM pathway plays a crucial role in pollen viability and reproductive success in plants and several mutants in this pathway, such as *nrpd1a, clsy3*, several AGO genes exhibit pollen sterility (Wang et al. 2020; Hari Sundar G et al. 2023; Pal et al. 2024). We performed pollen viability assay and found that the number of viable pollens was also low in osclsy4-kd, indicating possible reasons for reduced fertility (Supplemental Figure S1E). Similar to osclsy3-kd, the osclsy4-kd seeds were smaller than wild type PB1 seeds (Fig. 1D). However, osclsy4-kd seeds were wider than PB1 and osclsy3-kd seeds (Fig. 1D, E). While both clsy3-kd and clsy4-kd led to smaller seeds, grain chalkiness phenotype was observed only in osclsy4-kd endosperms (Fig. 1F and Supplemental Figure S1F). In monocots, endosperm plays a vital role not just in embryo development, but also in seed germination and hence major defects in endosperm results in germination defects (Yan et al. 2014). In seed germination assays, we observed slow germination of osclsy4-kd plants (Fig. 1G). The emergence and growth of coleoptiles were delayed in osclsy4-kd when compared to control seeds. Our results suggest that OsCLSY4 is also an important regulator of rice reproduction and endosperm development.

**Figure 1.**
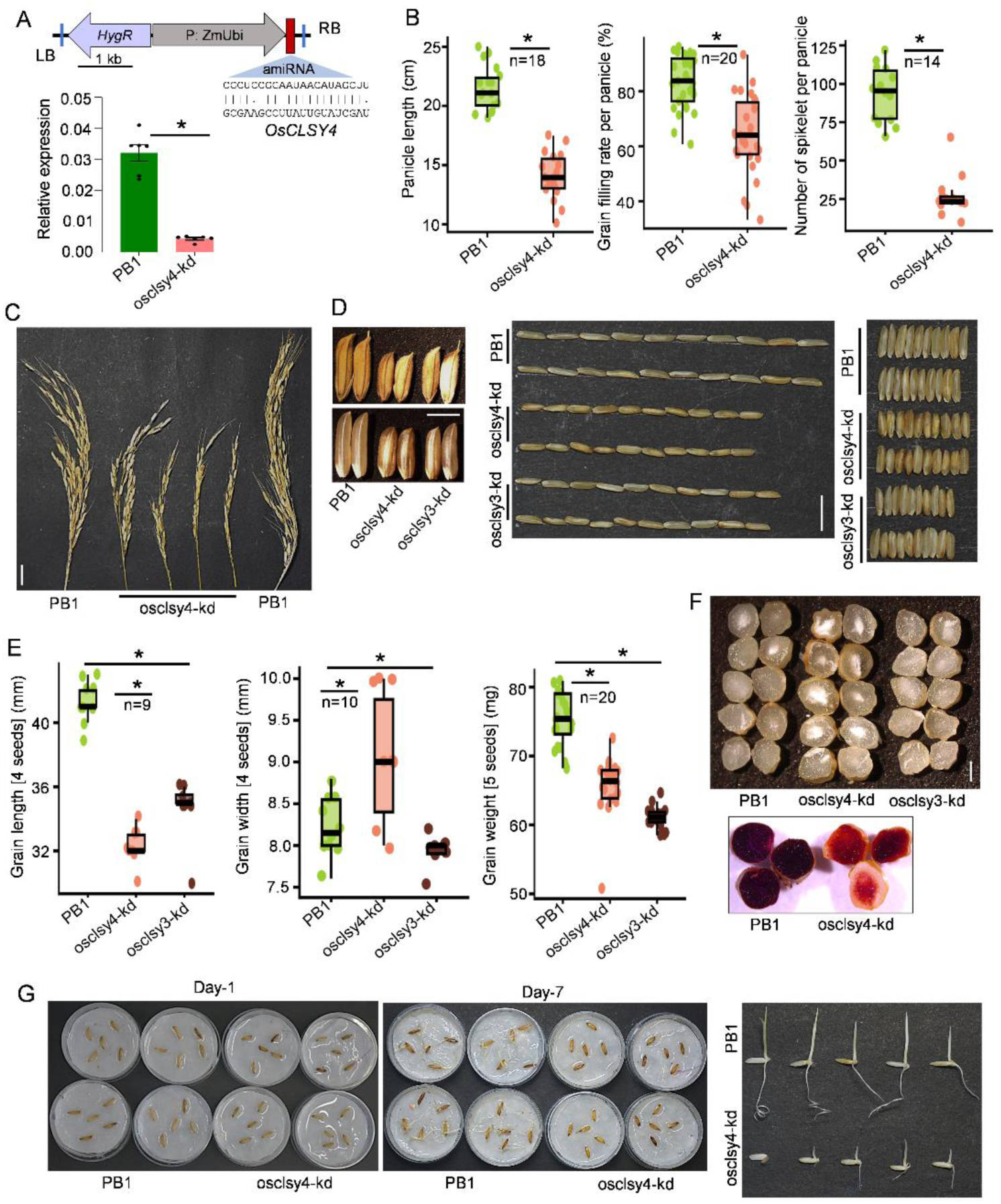
Endosperm-related phenotypes of osclsy4-kd plants. (A) T-DNA vector map of AmiR construct. AmiR is driven by maize Ubi promoter. RB-right border, LB-left border. Barplot showing levels of *OsCLSY4* expression in clsy4-kd endosperm (EN) is shown. *OsActin* served as internal control. Error bar-SE. *-significant. Two-tailed Student’s *t*-test. (B) Boxplots showing panicle phenotypes. In box plots, *-significant (two-tailed Student’s *t*-test). (C) Images showing panicle morphology. SB (Scale Bar)-2 cm. (D) Images showing morphology of osclsy3-kd and osclsy4-kd seeds. SB-1 mm. Length (middle panel) and width (right) of 10 seeds across lines are shown. (E) Boxplots showing seed length, width and weight in osclsy4-kd and osclsy3-kd. (F) Seed­phenotype of osclsy4-kd and osclsy3-kd dry grains (top) and iodine-staining (bottom) of osclsy4-kd endosperm sections. SB-0.5 mm. (G) Germination-phenotype of osclsy4-kd. Seeds were germinated and photographed. Individual germinating seeds after 7-days in water (Left) and 5-days in media (Right) are shown, respectively.

### OsCLSY4 regulated a large number of sRNA loci in rice endosperm

In *Arabidopsis,* CLSYs recruit Pol IV at specific TEs and repeat regions to initiate silencing. The Pol IV generates transcripts that are processed into 24 nt sRNAs from TE and repeat regions and these siRNAs induce DNA methylation (Zhou et al. 2018). To check the role of OsCLSY4 in regulating the accumulation of sRNAs in rice endosperm, we performed sRNA sequencing in 20 days-old clsy4-kd endosperm tissues. More than 90% of an average of 25 million reads mapped to the rice genome (Supplemental Table 1). As expected, we found drastic reduction of 24 nt sRNAs in osclsy4-kd endosperm-derived tissues similar to that of clsy3-kd lines (Fig. 2A). To compare the importance of both OsCLSY3 and OsCLSY4 in endosperm, we counted 20-25 nt sRNAs in both genotypes (Supplemental Data 1 and Supplemental Data 2). The analysis indicated that 23-24 nt size class sRNAs were more drastically reduced in osclsy4-kd when compared to osclsy3-kd, indicating that the majority of 24-nt sRNAs accumulating in endosperm are under OsCLSY4 control (Supplemental Figure S2A). Also, in osclsy4-kd, we observed 1500 OsCLSY4-dependent gained sRNA loci (Supplemental Figure S3A). Among the pools of 21, 22 and 24 nt sRNAs, sRNAs that had 5’ A were specifically reduced in osclsy4-kd, indicating their possible association with AGO4 clade proteins in wild type conditions (Supplemental Figure S2B). To find OsCLSY4-dependent 23-24 nt sRNAs, we performed ShortStack based analysis and found that the expression of more than 70% of 23-24 nt sRNA loci (10521 CLSY4-dependent loci) that are expressed more than 5 rpm in PB1 were reduced in osclsy4-kd endosperm tissues (Fig. 2B). These sRNAs originated from class I (TEs that replicate in genome by copy paste mechanism) and class II (TEs that replicate *via* cut paste mechanism) TEs as well as genic regions (Fig. 2C). In agreement with their OsCLSY4 dependency, these sRNA loci were reduced in osclsy4-kd (Fig. 2D). On the other hand, 21-22 nt sRNAs from miRNA loci were unaltered in osclsy4-kd endosperm, indicating that CLSY4 majorly regulated sRNAs associated with RdDM (Fig. 2E). Major families of TEs such as Copia, LINE, LTRs, MITEs and other repeat-derived 23-24 nt sRNAs were significantly reduced in osclsy4-kd lines (Fig. 2F and Supplemental Figure S2C). We found that many of the OsCLSY4-dependent 23-24 nt sRNAs from TEs overlapped with OsCLSY3-dependent loci 23-24 nt loci that clearly showed redundant function of both OsCLSYs in TEs (Fig. 2G). We also observed a reduction in a large number of sRNAs, particularly in individual CLSY kd lines which suggested their non-redundant function (Fig. 2H and Supplemental Figure S2D). All these results collectively suggested that OsCLSY4 is an important RdDM regulator in rice endosperm.

**Figure 2.**
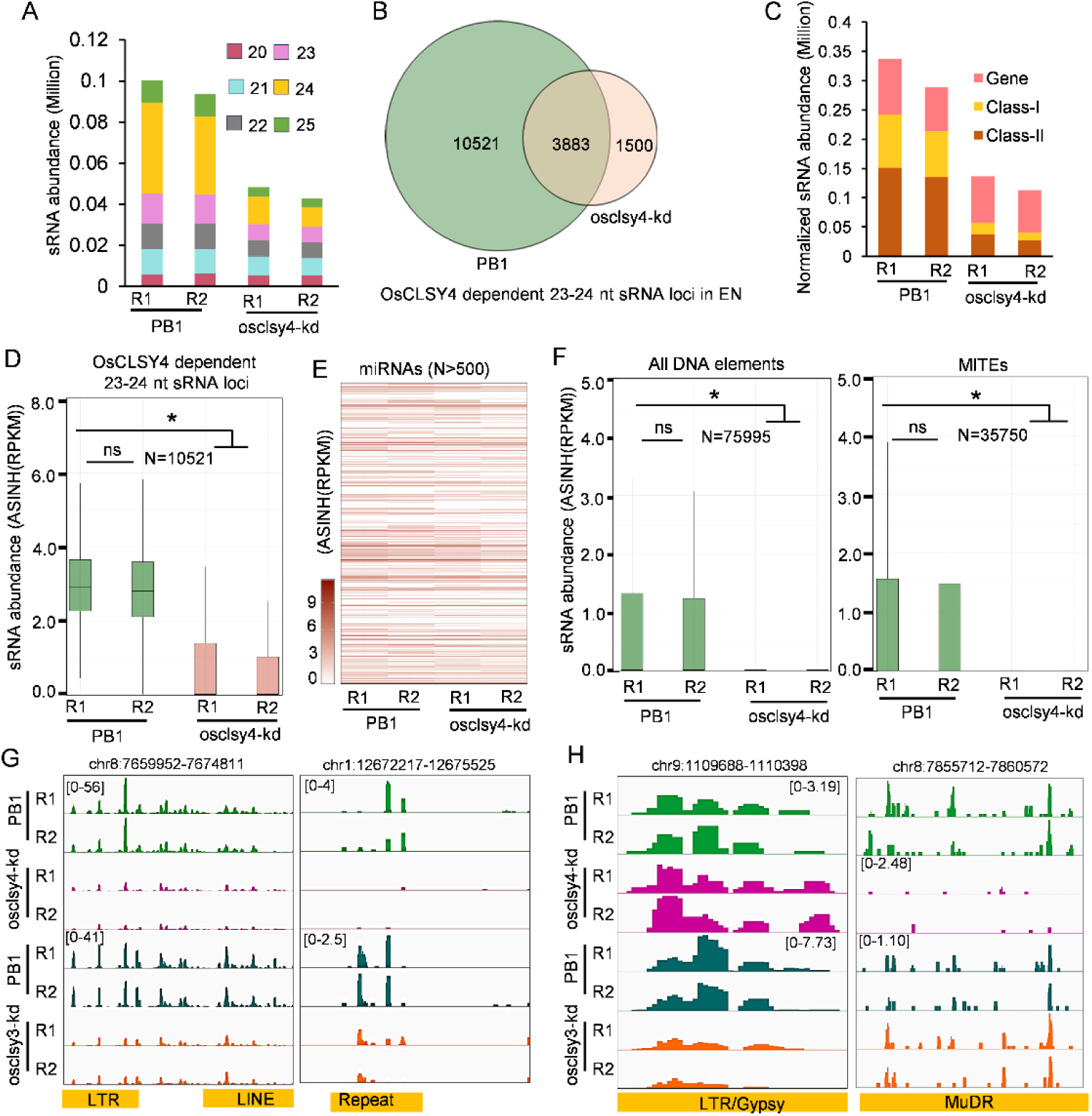
Endosperm sRNAs are globally reduced in osclsy-kd lines. (A) Stacked barplot showing abundance of sRNAs (20-25 nt) in PB1 and osclsy4-kd EN tissues. (B) Venn diagram showing 23-24 nt sRNA loci across PB1 and osclsy4-kd EN. (C) Plot showing 23-24 nt sRNA abundance across different genomic features. (D) Boxplot showing abundance of OsCLSY4-dependent 23-24 nt sRNA loci in PB1 and osclsy4-kd EN. *-significant, ns-non-significant (Wilcoxon test p < 0.01). (E) Heatmap showing expression of miRNAs in PB1 and clsy4-kd EN. ASINH converted RPKM values were used for the heat map. (F) Boxplots showing abundance of 23-24 nt sRNAs in different TE types. N-number of loci. (G) and (H) IGV screenshots showing expression of 23-24 nt OsCLSY3- and OsCLSY4-dependent shared (left) and unique sRNA loci (right).

### OsCLSY4 and OsCLSY3 non-redundantly regulate accumulation of sRNAs in endosperm

In *Arabidopsis*, CLSY proteins function both non-redundantly and redundantly (Zhou et al. 2018; Martins and Law 2023). AtCLSYs show a tissue- and locus-specific regulation along with shared loci (Zhou et al. 2018, 2022). In order to find if OsCLSY3 and OsCLSY4 act on specific loci in endosperm tissue, we analyzed published sRNA datasets derived from osclsy3-kd endosperm tissues (Supplemental Data 2 and Supplemental Data 3). In osclsy3-kd, we found more than 60% of 23-24 nt sRNA loci reduced when compared to WT control (Supplemental Table 2). Along with these OsCLSY3-dependent 23-24 nt sRNA loci, we also obtained 583 sRNA loci that were upregulated in osclsy3-kd (Supplemental Figure S3B, C). Upon overlapping OsCLSY3 and OsCLSY4-dependent 23-24 nt sRNA loci, we observed 1014 sRNA loci that overlapped with each other (‘shared loci), while 9507 OsCLSY4-dependent sRNA loci (henceforth, ‘OsCLSY4-unique loci’) and 3889 CLSY3-dependent loci did not overlap with the other CLSY (henceforth, ‘OsCLSY3-unique loci’) (Fig. 3A).

**Figure 3.**
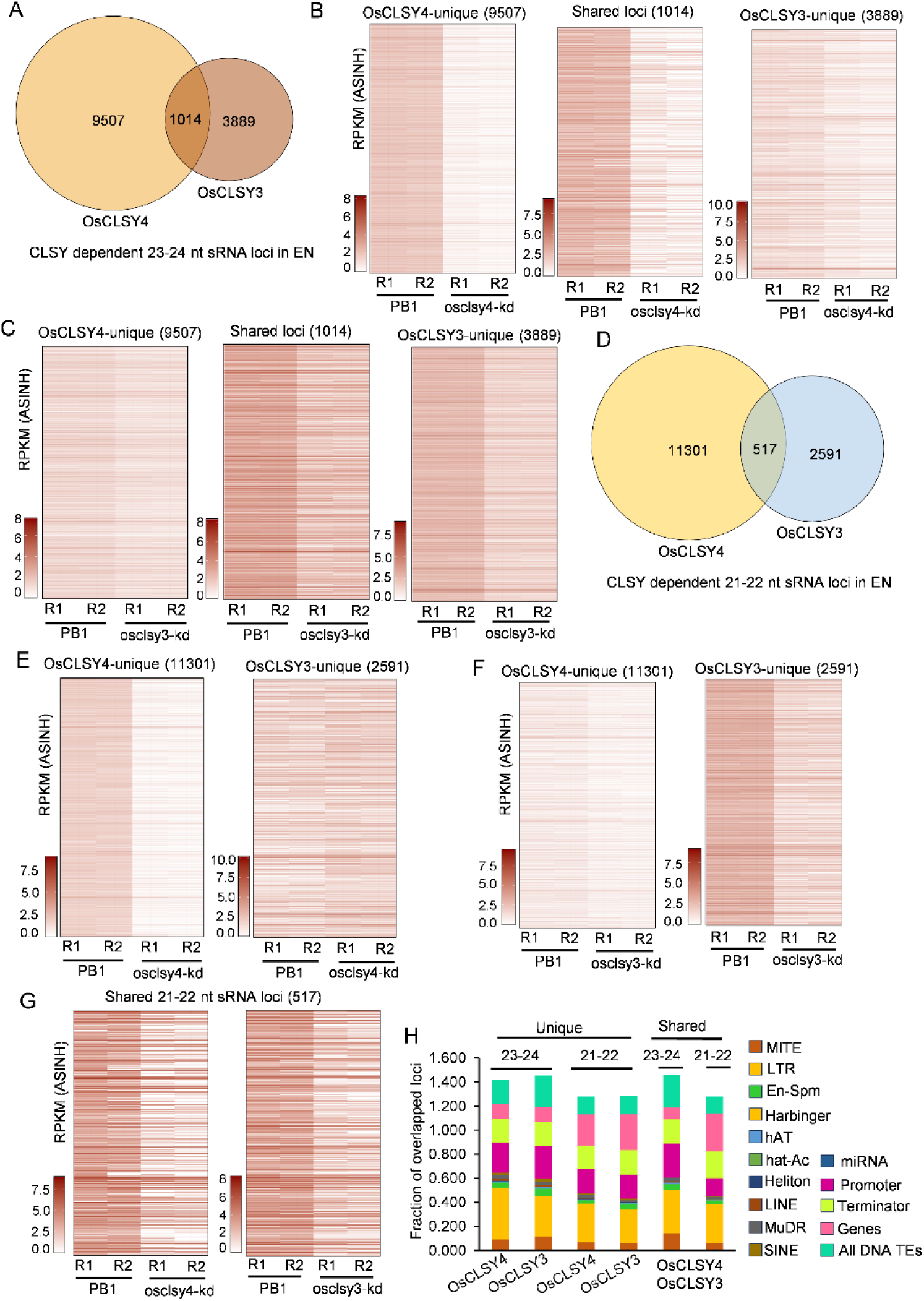
CLSYs regulate endosperm expressing sRNAs in non-redundant ways. (A) Venn diagram representing overlap between OsCLSY3 and OsCLSY4-dependent 23-24 nt sRNA loci. (B), (C) Heatmaps showing 23-24 nt sRNA abundance across OsCLSY4-unique, OsCLSY3-unique and shared sRNA loci. ASINH converted RPKM values were used for heatmaps. (D) Venn diagram representing overlap between OsCLSY3 and OsCLSY4-dependent21-22 nt sRNA loci. (E),(F),(G) Heatmaps showing 23-24 nt sRNA abundance across OsCLSY4-unique, OsCLSY3-unique and shared sRNA loci, respectively. (H) Stacked bar plots representing overlap of OsCLSY3- and OsCLSY4-unique as well as shared sRNA loci with different genomic features.

As expected, OsCLSY4-unique loci were much more reduced in abundance in osclsy4-kd while OsCLSY3-unique loci were more reduced in osclsy3-kd (Fig. 3B, C and Supplemental Data 3). We also observed a marginal reduction of OsCLSY4-unique and OsCLSY3-unique sRNAs in osclsy3-kd and osclsy4-kd endosperm tissues, respectively (Fig. 3B, C) In order to find out how prevalent shared loci are across TE and other repeat features, we extended the sRNA loci width to 1 kb on both sides and identified a total of 1964 loci that were possibly redundantly regulated by both CLSYs in endosperm tissues (Supplemental Figure S3D). All the results collectively suggested that rice CLSYs function largely in a non-redundant fashion in endosperm tissues.

Similar to *Arabidopsis*, along with predominant 24 nt sRNAs, there are also 21-22 nt sRNAs in rice endosperm (Erdmann et al. 2017; Pal et al. 2024). A majority of the 21-22 nt sRNA loci also overlapped with 23-24 nt sRNA loci, indicating that their precursors are targeted by multiple DCLs (Erdmann et al. 2017). ShortStack analysis identified 11818 CLSY4-dependent 21-22 nt sRNA loci in rice endosperm (Supplemental Figure S3E-F). In this analysis, we also identified around 2300 sRNA gained loci (Supplemental Figure S3E). Around 3987 sRNA loci showed downregulation of both 21-22 and 23-24 nt sRNAs in osclsy4-kd endosperm indicating that OsCLSY4 is upstream of sRNA loci having both 23-24 and 21-22 size sRNAs (Supplemental Figure S3G). Similarly, Some OsCLSY3-dependent sRNA loci were also of both size classes (Supplemental Figure S3H-J).

Similar to 23-24 nt sRNAs, there were 21-22 nt sRNAs exclusively regulated by both OsCLSY4 and OsCLSY3. We identified 11301 OsCLSY4-unique and 2591 CLSY3-unique 21-22 nt sRNA loci (Fig. 3D). As expected, OsCLSY4-unique sRNA loci were downregulated in osclsy4-kd but not in osclsy3-kd (Fig. 3E). OsCLSY3-unique sRNA loci were mostly downregulated in osclsy3-kd (Fig. 3F). The shared 517 loci were reduced in both kd lines as expected (Fig. 3G). Upon extending the length of 21-22 nt sRNA loci, we obtained around 2047 loci in which OsCLSY3 and OsCLSY4 might be functioning redundantly in the endosperm (Supplemental Figure S3K).

In order to identify if there is a preference for a genomic feature among CLSYs, we overlapped unique as well as shared sRNA loci with different TEs and other genomic features. We did not observe any preference for a specific feature or TE (Fig. 3H).

Even among 21-22 nt sRNAs that are majorly derived from genic regions, both CLSYs did not show any preference (Fig. 3H).

### OsCLSY4 regulates site-specific DNA methylation in endosperm

CLSYs regulate DNA methylation through the RdDM pathway in both vegetative and reproductive tissues across dicots and monocots (Yang et al. 2018; Zhou et al. 2018). Since OsCLSY4 is also expressed in leaf tissues, we performed targeted BS-PCR to check the DNA methylation level in leaves of osclsy4-kd lines. The 5S-rDNA repeat was specifically hypomethylated in osclsy4-kd leaves conclusively indicating that OsCLSY4, but not endosperm-specific OsCLSY3, regulates RdDM in vegetative tissues (Supplemental Figure S4).

To study the role of OsCLSY4 in the regulation of methylation, we performed whole genome bisulfite sequencing of genomic DNA derived from 20-day-old endosperm tissues (Supplemental Figure S5A). We identified a total of 27345 differentially methylated regions (DMR) in osclsy4-kd in comparison to 27022 DMRs in osclsy3-kd endosperm. In osclsy4-kd, a total of 5984 CG, 10365 CHG and 10997 CHH hypomethylated loci were identified (Supplemental Data 4). These numbers were comparable to OsCLSY3-dependent DMRs that we identified previously, indicating that OsCLSY4 also contributes to RdDM in endosperm (Supplemental Data 5). Interestingly, we found that 9904 DMRs were common between osclsy4-kd and osclsy3-kd lines (Fig. 4A). Among these shared DMRs, a total of 3626 were exclusively CHH DMRs (Fig. 4B). As expected, DNA methylation was reduced in OsCLSY3- and OsCLSY4-DMRs in respective kd lines. However, OsCLSY3 and OsCLSY4-DMRs also exhibited significant reduction in osclsy4-kd and osclsy3-kd lines, indicating that there is redundancy in the activities of these CLSYs in inducing RdDM at some loci (Fig. 4C, 4D). OsCLSY4-dependent sRNA loci overlapped with OsCLSY4-dependent DMRs indicating that OsCLSY4 regulates DNA methylation at these sites (Fig. 4E). Among shared loci, DNA methylation at CHH sites were reduced in both kd lines as expected (Fig. 4F). DNA hypomethylation at several TEs loci, especially MITEs, were observed in osclsy4-kd lines correlating with sRNA reduction (Fig. 4E, F). These results indicate that, along with OsCLSY3, OsCLSY4 also regulates site-specific DNA methylation in rice endosperm.

**Figure 4.**
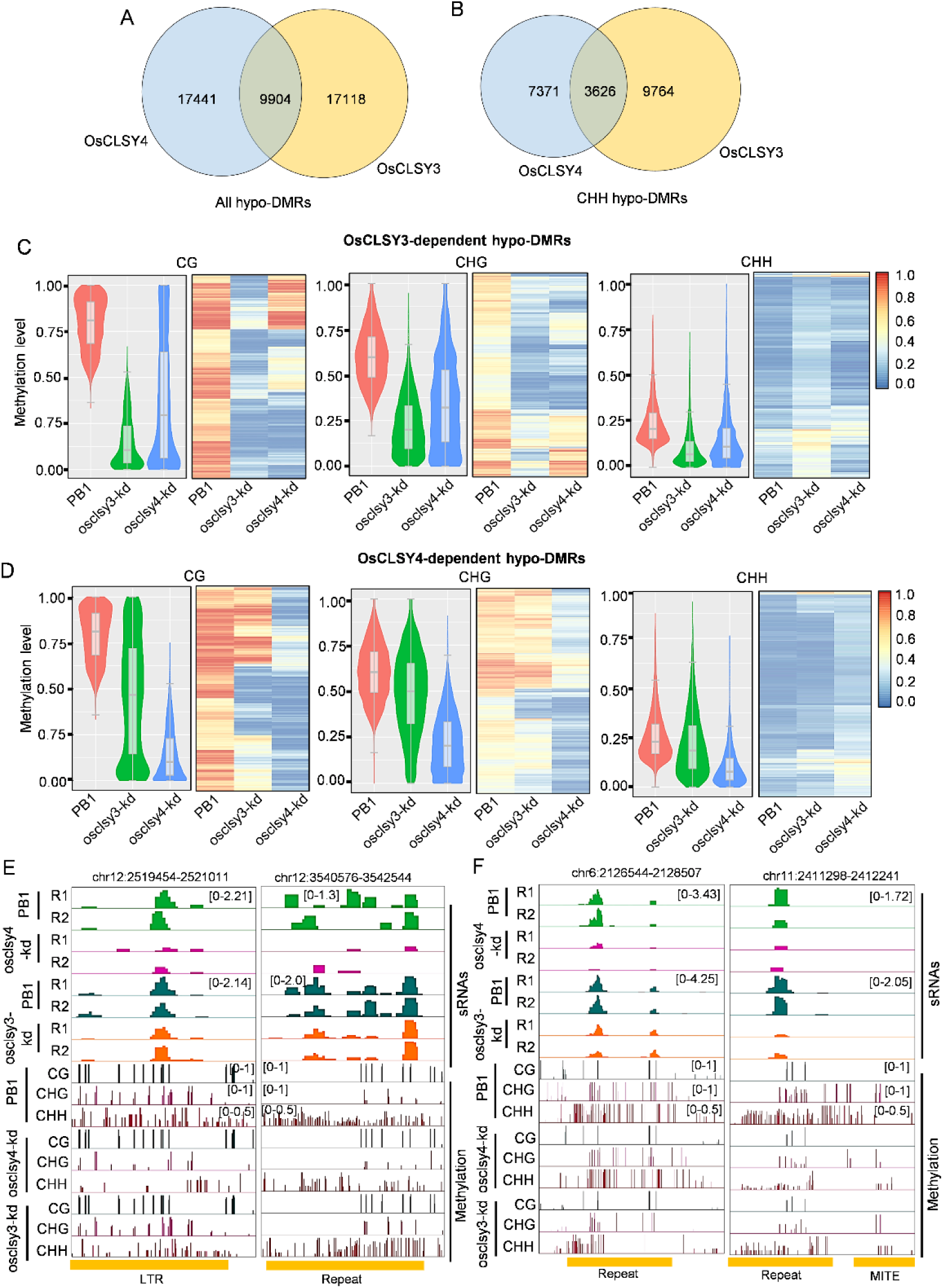
Both OsCLSY3 and OsCLSY4 regulate methylation through RdDM pathway in endosperm. (A), (B) Venn-diagrams showing overlap between all hypo-DMRs and CHH context hypo-DMRs in both kd lines in EN, respectively. (C), (D) Violin-plots and heatmaps showing OsCLSY3 and OsCLSY4-dependent hypo-DMRs, respectively. (E), (F) IGV screenshots representing sRNA and DNA methylation status of CLSY4-unique and shared loci, respectively.

### osclsy4-kd plants exhibit pronounced hypermethylation across repeats and genic regions

Previous studies in *Arabidopsis* as well as in rice indicated pronounced hypermethylation at several genomic loci when different CLSY genes were mutated (Yang et al. 2018; Zhou et al. 2018; Martins and Law 2023; Pal et al. 2024). However, the regions showing hypermethylation were neither shared loci between individual CLSYs, nor they were under the direct control of another CLSY member. While the mechanism of this hypermethylation is unknown, it is possible that more than one factor contributed to this phenomenon. Similar to osclsy3-kd, we observed pronounced hypermethylation at several genomic loci specifically at CHH sites in osclsy4-kd (Supplemental Figure S5A-B). In osclsy3-kd, 1976 CG,7930 CHG and 34070 CHH hypermethylated loci were identified using DMRcaller. About 1062 CG, 2617 CHG and 22761 CHH hypermethylated loci were identified in osclsy4-kd lines (Supplemental Data 4, Supplemental Data 5 and Supplemental Figure S5C). We overlapped hyper-DMRs between the OsCLSY3 and OsCLSY4 and identified 10021 shared hyper-DMRs across all sites and 7855 shared CHH hyper DMRs (Supplemental Figure S5A). To identify possible genomic preference between hypo-and hyper-DMRs of both kd lines, we overlapped the DMRs with different genomic features and identified that the DMRs arose from all genomic features without any specific preference (Supplemental Figure S5B).

In our analysis, different categories of TEs, such as MITEs, Gypsy and SINE elements showed DNA hypomethylation mainly in CHG context as expected. However, there were many TEs having hypermethylation, mainly in CHH context (Fig. 5A). Among hypermethylated sites, abundance of 21-22 nt sRNAs were not observed in any of the lines, indicating that CHH methylation did not arise from non-canonical 21-22 nt sRNA-mediated DNA methylation (Fig. 5B-D). Surprisingly, a majority of the hypermethylated loci were not overlapping with any class of sRNAs (Fig. 5B-D). In order to understand if osclsy4-kd led to altered expression of components of DNA methylation/demethylation machinery, we tested expression levels of DNA demethylases and methyltransferases and identified that two ROS genes, namely OsROS1a (Os01g0218032) and OsDML4 (Os03g0110800) were downregulated in osclsy4-kd (Supplemental Data 7). Interestingly, the expression of DRM2 was also significantly reduced in osclsy4-kd endosperm. These results collectively indicate that CLSY proteins might be suppressing DNA methylation and/or DNA demethylation at several sites to result in DNA hypermethylation in rice clsy mutants. It will be interesting to identify whether these alterations are cell-type/tissue-specific and/or developmentally regulated.

**Figure 5.**
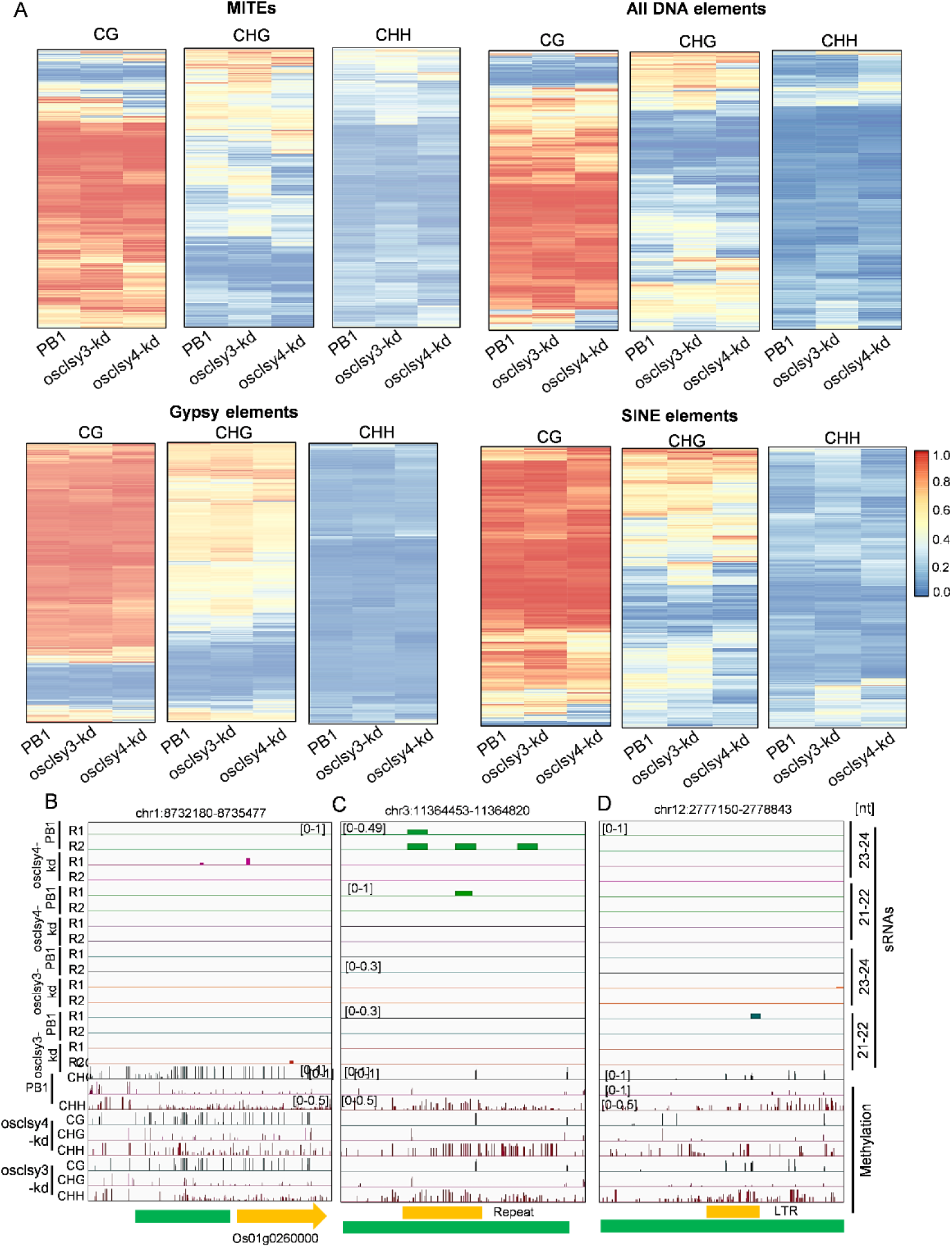
sRNA independent hypermethylation in osclsy3-kd and osclsy4-kd EN tissues. (A) Heatmaps showing DNA methylation levels across different TEs in kd lines of both CLSYs in EN. (B), (C),(D) IGV screenshots showing hyper methylated OsCLSY4, shared and OsCLSY3 loci in EN, respectively.

### OsCLSY3 and OsCLSY4 redundantly regulate siren loci in rice

In endosperm tissues, a unique set of sRNA loci called siren contribute to most of the sRNAs derived from fewer loci and regulate specific genes involved in its development (Rodrigues et al. 2013, 2021; Grover et al. 2018, 2020; Burgess et al. 2022; Dziasek et al. 2024). They predominantly generate from the maternal tissues (Grover et al. 2018; Burgess et al. 2022; Dziasek et al. 2024). Siren sRNAs act as hybridization barriers by regulating parental dose of gene expression (Dziasek et al. 2024). In *Arabidopsis* and *Brassica rapa*, expression of siren sRNAs are regulated by reproductive-tissue specific CLSY3 and CLSY4 (Grover et al. 2020; Burgess et al. 2022; Zhou et al. 2022). We explored if OsCLSY4 is involved in the regulation of siren expression in rice endosperm similar to OsCLSY3 (Pal et al. 2024). Among the 797 siren loci in rice, most were reduced in abundance in osclsy4-kd endosperm (Supplemental Data 6). These loci were also reduced in osclsy3-kd lines indicating that both these CLSYs regulate expression of siren loci redundantly (Fig. 6A-C). As observed in *Arabidopsis* (Dziasek et al. 2024), several siren loci were heavily methylated and the status of DNA methylation remained unchanged in osclsy3-kd and osclsy4-kd lines (Fig. 6D), however, in selected siren loci, we also observed DNA hypomethylation in both kd genotypes (Fig. 6B). A total of 38 siren adjacent genes, such as Os02g0527200, Os10g0177200 and Os01g0629900 were significantly upregulated in kd lines. Also, around 70 genes were downregulated in osclsy4-kd (Fig. 6E). Although, the importance of DNA methylation and its contribution to siren RNA loci expression is unknown, it is clear that both OsCLSYs play a major role in their regulation and indirectly the expression of genes associated with them. It is important to note that the homologs of rice CLSY3 and CLSY4, i.e., AtCLSY3 and AtCLSY4, also regulate siren loci expression in ovules and endosperm indicating possible conserved functions of this clade of CLSY members.

**Figure 6.**
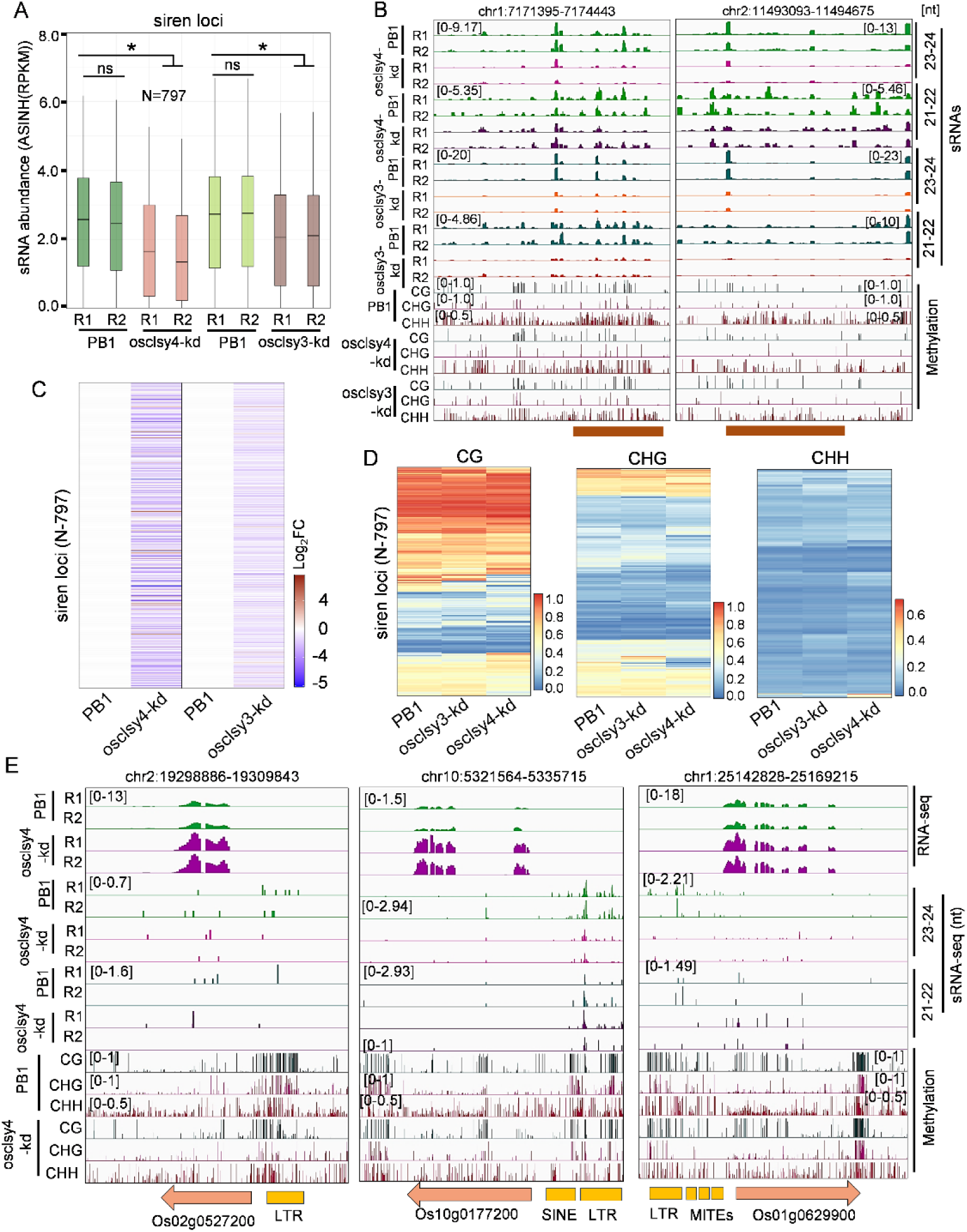
Both CLSYs redundantly regulate siren loci in rice. (A) Boxplot showing expression of 23-24 nt sRNAs in PB1, and osclsy3-kd and osclsy4-kd EN. *-significant, ns-non-significant (Wilcoxon test p < 0.01). (B) IGV screenshots depicting expression of 23-24 and 21-22 nt sRNAs from siren loci. (C) Heatmaps showing expression of siren loci (N-797) in the osclsy4-kd and osclsy3-kd EN tissues. (D) Heatmaps showing DNA methylation level in siren loci in kd lines EN. (E) IGV screenshots showing expression of siren loci adjacent genes in osclsy4-kd EN.

### OsCLSY4-dependent sRNA loci control expression of neighboring genes through RdDM

In rice, similar to osclsy3-kd, osclsy4-kd lines also displayed endosperm defects. To understand the molecular basis for the chalkiness phenotype observed in osclsy4-kd, we performed transcriptome analysis of 20-day-old osclsy4-kd endosperms. We documented absence of green-tissue specific marker gene expression indicating that the tissues taken for analysis were free from maternal-tissue contamination (Supplemental Figure S6A). A total of 1456 upregulated and 1875 downregulated genes were observed in osclsy4-kd (Fig. 7A). Although specific pathway genes were not altered, several metabolism-related genes were mis-expressed in osclsy4-kd lines (Supplemental Figure S6B). Since osclsy4-kd but not osclsy3-kd endosperm showed chalkiness phenotypes indicating a lack of progression to cellularization, we overlapped differentially expressed genes (DEGs) between both kd lines. We identified 2206 and 1628 unique DEGs in osclsy4-kd and osclsy3-kd endosperms, respectively, indicating that both CLSYs play unique roles in endosperm development (Fig. 7B). Expression of genes such as Os01g0726250 and Os08g0127900 (an endosperm-specific gene 115/ Similar to Globulin 1) behaved similarly between both kd lines (Fig. 7C). We identified 30 upregulated and 31 downregulated genes shared between both kd lines (Supplemental Figure S6C). There were also genes that showed clear differences between both kd lines. Genes such as Os07g0568700 (FORAL ORGAN REGULATOR 1) and Os12g0268000 (Cytochrome P450 monooxygenase) had opposite expression patterns between osclsy3-kd and osclsy4-kd lines (Supplemental Figure S6D). Some of the gene expression abnormalities might be due to vegetative and reproductive defects observed in osclsy4-kd.

**Figure 7.**
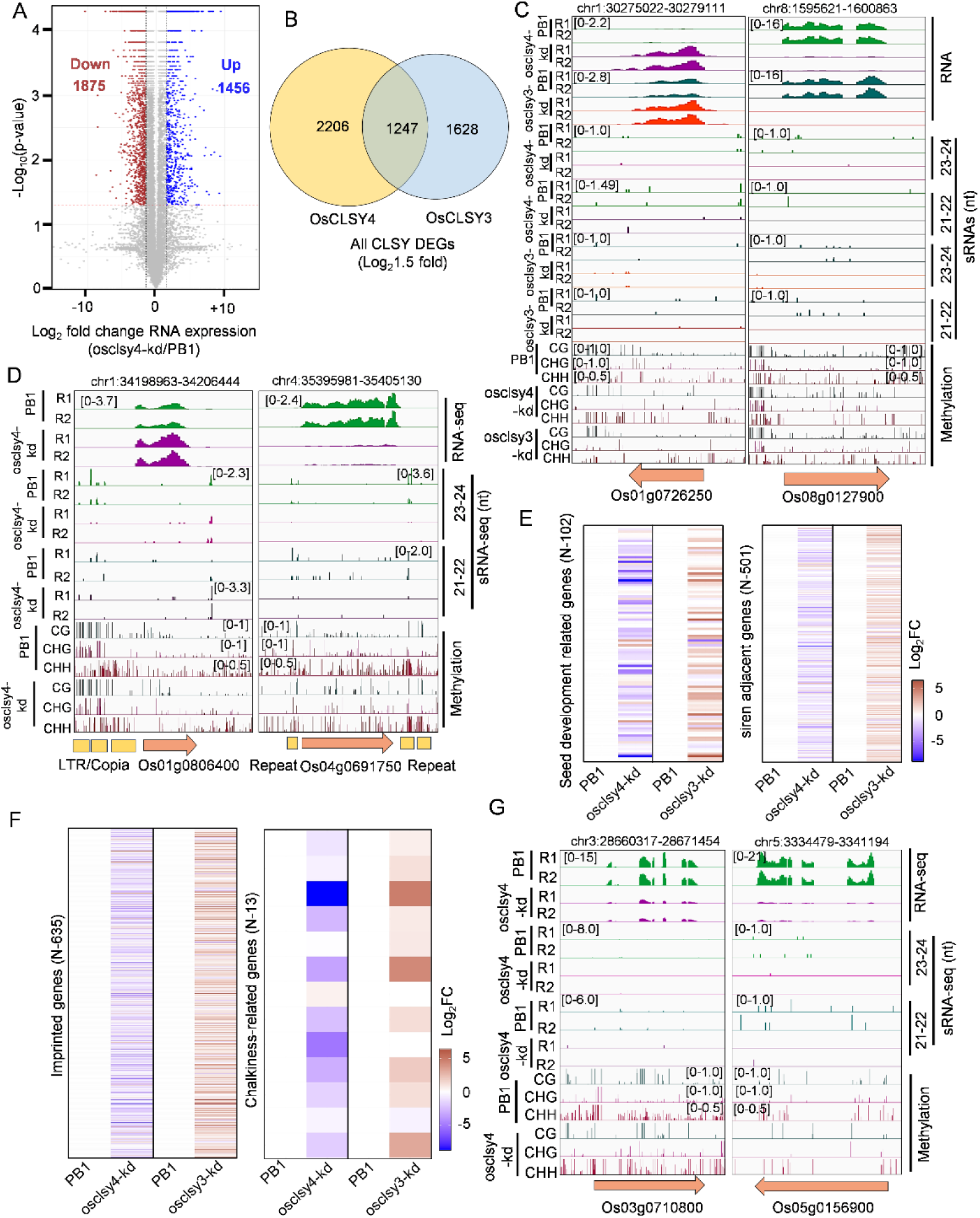
OsCLSY4 regulates expression of many crucial endosperm-specific genes in rice. (A) Volcano plot showing expression of genes in clsy4-kd EN. (B) Venn diagram representing overlap between DEGs (Log_2_fold change-1.5) between osclsy3-kd and osclsy4-kd EN. (C) IGV screen shots showing expression of upregulated and downregulated genes in osclsy4-kd and osclsy3-kd EN. (D) IGV screenshots showing upregulated and downregulated genes in osclsy4-kd EN. (E) and (F) Heatmaps showing expression of seed-development related, siren adjacent, imprinted and chalkiness-related genes in osclsy3-kd and osclsy4-kd EN tissues, respectively. Log_2_fold change (FC) RPKM values were used for heatmaps. (G) IGV screenshots showing CLSY4-mediated regulation of two chalkiness-related genes in osclsy4-kd EN.

**Figure 8.**
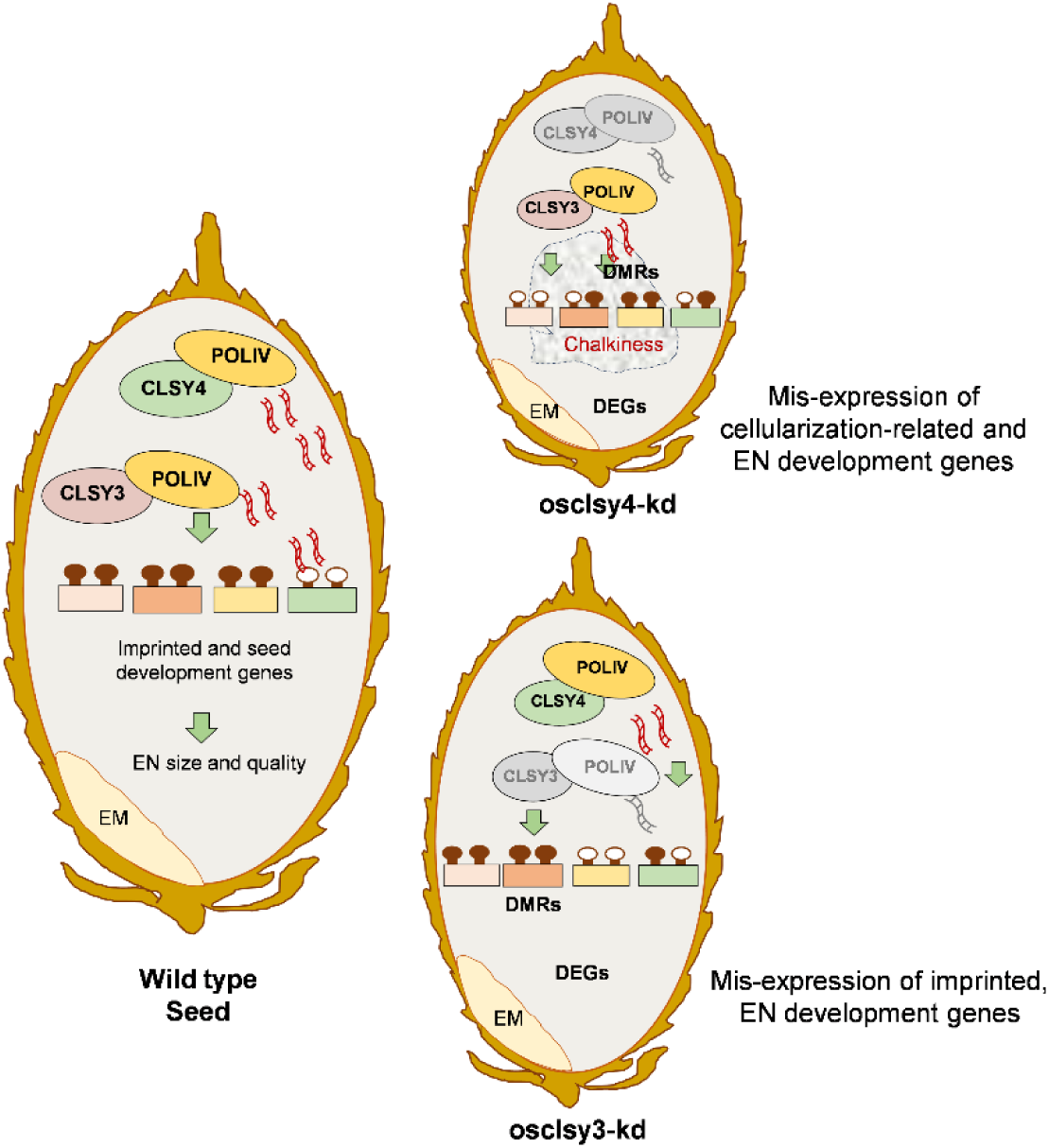
OsCLSY4 is also a crucial regulator of rice endosperm development. OsCLSY4 and OsCLSY4 both are important for proper endosperm development in rice. Both the kd lines showed smaller endosperm but only osclsy4-kd showed grain-chalkiness phenotype. Both rice CLSYs control many unique loci as well as a few shared loci in EN. Those sRNAs direct DNA methylation (filled lollypop) and induced associated epigenetic regulations that are crucial for proper endosperm development. Both kd lines show hypomethylation at their dependent loci (unfilled lollypops). EM-embryo, EN-endosperm, DEGs-differentially expressed genes, DMRs-differentially methylated regions.

To explore the role of OsCLSY4-associted RdDM pathway in controlling expression of genes, we checked the sRNA and methylation levels in the vicinity of upregulated DEGs. The genes such as Os01g0806400 (DUF617) and Os01g0136100 (OsHsp16.9A) showed clear upregulation in osclsy4-kd (Fig. 7D and Supplemental Figure S6E). These gene promoters contain TEs and repeats under the control of OsCLSY4. Several seed development-associated genes were downregulated in osclsy4-kd mainly due to DNA hypermethylation at their promoters. For example, Os11g054650 (glycosyltransferase 6) and Os04g06917500 (phosphoribosylanthranilate transferase) were downregulated in osclsy4-kd endosperm and their promoters or terminator were hypermethylated (Fig. 7D and Supplemental Figure S6E). These results collectively suggested that OsCLSY4 regulates expression of many genes through RdDM pathway. Since many genes were uniquely mis-expressed between osclsy3-kd and osclsy4-kd endosperm, it is clear that they regulate endosperm development likely in different stages/tissue/cell types.

### OsCLSY4 has distinct targets in vegetative tissues and endosperm

Unlike OsCLSY3, OsCLSY4 is expressed in both vegetative and reproductive tissues in rice. In leaf tissues, OsCLSY4 is the major CLSY member and it appears to target genomic regions such as TEs and repeats (Xu et al. 2023). To understand if the sRNA loci regulated in vegetative tissue and endosperm originate from the same regions, we analyzed and compared OsCLSY4-dependent sRNA loci across these tissues (Supplemental Data 8). Similar to endosperm tissue, 23-24 nt sRNAs were also reduced in class I, class-II TEs and genes (Supplemental Figure S7A). Interestingly, there were 3544 CLSY4-dependent 23-24 nt sRNA loci overlapping between these tissues. However, a majority of the OsCLSY4-dependent sRNA loci were unique between leaf (69721) and endosperm (6977) tissues, indicating a tissue-specific preference of CLSY4 for genomic loci (Supplemental Figure S7B). OsCLSY4-dependent sRNA loci originated from TEs and genes in a tissue-specific manner (Supplemental Figure S7C-S7F). This altered preference of OsCLSY4 might be due to variation in its expression level, possibility of its distinct protein interacting partners or other epigenetic marks. Since CLSY4-dependent sRNAs were mostly different between leaves and endosperm, it is possible that a different set of genes are under its control in these tissues. In agreement with this, several transcription factors specifying the recruitment of AtCLSY3 to initiate RdDM were recently identified, and they had specific roles in male and female reproductive tissues (Wu et al. 2025; Xu et al. 2025). To explore genes that are regulated by OsCLSY4 in two tissues, we compared the DEGs and found only 119 genes that were shared between two tissues (Supplemental Figure S7G and Supplemental Data 9). All these results collectively suggested a tissue-specific role for ubiquitously expressed OsCLSY4, contributing to tissue-specific sRNA expression and RdDM.

### OsCLSY4 regulates expression of many seed development and chalkiness-related genes in rice

The triploid tissue of endosperm expresses a unique set of genes that are important for the regulation of development, including nutrient accumulation. Endosperm development and growth is a complex process regulated by several signaling, hormonal and epigenetic pathways (Li et al. 2019, 2022). We observed the mis-expression of many seed development-related genes connected to these pathways in endosperms of both the kd lines (Fig. 7E, Supplemental Figure S6E-S6F). Around 15 seed development-related genes were among the DEGs in osclsy4-kd. Genes such as OsBZR1, OsAGO2, GW6, OsYUCCA11 and NF-YC12 (Abu-Zaitoon et al. 2012; E et al. 2018; Gao et al. 2019; Li et al. 2022), which are well-known regulators of endosperm development in rice, were significantly downregulated in osclsy4-kd. Key metabolism-associated genes such as OsGAD2, AWN1 and RAG2 were significantly upregulated in the clsy4-kd lines. AWN1 (Os04g0350700) is a negative regulator of seed size and its reduced expression correlated with shorter seeds in osclsy4-kd line (Wang et al. 2019). Among the MADS-box genes that regulate floral development, cellularization and developmental programmed cell death in rice endosperm (Tao et al. 2018; Kong et al. 2022; Doll and Nowack 2024), we observed total of 7 MADS-box genes -OsMADS3, OsMADS4, OsMADS6, OsMADS8, OsMADS17, OsMADS18 and OsMADS29 that were significantly downregulated in osclsy4-kd (Supplemental Figure S6F). The siren loci adjacent genes were also mis-expressed in osclsy4-kd lines (Fig. 7E). Among the 635 imprinted genes in rice, 139 imprinted genes were significantly mis-expressed in osclsy4-kd. Among them, 42 genes had a clear upregulation in osclsy4-kd (Fig. 7E and 7F). These results suggest that OsCLSY4 is also an essential and important epigenetic-regulator of gene expression in rice endosperm.

Interestingly, we observed grain-chalkiness phenotype in osclsy4-kd endosperm but not in osclsy3-kd (Fig. 1F). The grain-chalkiness is a complex phenotype which arises due to environmental stresses. Defects in starch, protein metabolism, transcription, organelle-development related genes lead to grain-chalkiness (Chen et al. 2024). Here, in osclsy4-kd, around 7 well-studied chalkiness-related genes such as OsNUDX7, OsMKK3, OsbHLH96, OsCIN2, GF14f, OsCrRLKL3 were significantly downregulated (Fig. 7G). The majority of those genes were unaltered upon osclsy3-kd. The two genes OsbHLH96, CHALK5 were also slightly upregulated in osclsy3-kd endosperm (Fig. 7F). All these results collectively suggested that the expression of upstream players of the RdDM pathway regulates chalkiness in rice.

## Discussion

The fate of endosperm is different in monocots when compared to dicots. In monocots such as rice, endosperm is retained and used during germination (Brown and Lemmon 2007; Liu et al. 2022). Epigenetic pathways such as DNA methylation, demethylation and histone modifications regulate endosperm development. Epigenetic players such as *OsFIE1, OsEMF2a, ZmFIE1* were identified as crucial players for genomic imprinting and endosperm development in monocots (Grossniklaus et al. 1998; Köhler et al. 2005; Luo et al. 2009; Zhang et al. 2012; Huang et al. 2016; Cheng et al. 2020, 2021; Tonosaki et al. 2021). Among flowering plants, RdDM also regulates parental dose balance to determine proper cellularization timing in endosperm tissues (Kirkbride et al. 2019; Dziasek et al. 2024). In most plants RdDM mutants exhibited severe reproductive abnormalities, such as pollen defect, low seed setting, seed size alterations, etc., clearly indicating the importance of RdDM pathway in seed or endosperm development process. How exactly the RdDM pathway recognizes TEs and repeats in a tissue-specific manner in rice is not well understood.

Among the three CLSYs, putative chromatin remodelers upstream of the RdDM pathway, OsCLSY3 and OsCLSY4 express in endosperm tissues. In endosperm, imprinted OsCLSY3 expresses at higher level than OsCLSY4, while OsCLSY1 is restricted to the embryo and early growth stages. OsCLSY3 majorly bound to LTR TEs as well as other genomic regions and targeted them for RdDM (Pal et al. 2024). This regulation was important for the expression of imprinted genes and seed development-associated genes. In this work, we show ubiquitously expressed OsCLSY4 also as a major component in rice endosperm development, targeting specific TEs and repeats different from those targeted by OsCLSY3. We found both osclsy3-kd and osclsy4-kd lines had smaller seeds but there were also many phenotypic differences in seeds between the kd lines.

It is important to delineate redundant and non-redundant functions of these CLSY proteins that promote endosperm development and imprinting in rice. As shown in *Arabidopsis*, we also observed OsCLSYs majorly act non-redundantly in endosperm but there were a large number of loci in which both function in a redundant manner. Both CLSYs act redundantly in the siren loci and associated genes. Interestingly, 23-24 nt sRNAs were much more reduced in osclsy4-kd than osclsy3-kd which further suggested that ubiquitously expressed OsCLSY4 might be the predominantly involved CLSY in rice, similar to AtCLSY1. Recent studies indicated that a large number of maternal sRNAs accumulated in the endosperm originated from surrounding maternal tissues (Grover et al. 2018, 2020; Dziasek et al. 2024). Unlike *OsCLSY3*, the *OsCLSY4* is expressed in the surrounding tissues in addition to endosperm and this CLSY might help in the accumulation of sRNAs in endosperm from surrounding maternal tissues. As reported in *Arabidopsis* endosperm (Erdmann et al. 2017), rice endosperm also produces a large number of 21-22 nt sRNAs that were under the control of both OsCLSY3 and OsCLSY4. Since the genes associated with siren loci are key regulators of imprinting, triploid block and post-zygotic reproductive barrier, CLSYs might act as crucial players in these processes. Sequence variation of OsCLSY3 between members of *Oryza* has been documented (Pal et al. 2024). Amino acid variations between both OsCLSY3 and OsCLSY4 between different rice accessions as well their expression variations might differentially regulate TEs and repeats thereby controlling key agronomically important traits. In *Arabidopsis*, CLSY1 D538E mutation alteration is involved in the regulation of lateral root numbers specifically in low K^+^ conditions (Shahzad et al. 2020).

DNA hypermethylation at specific regions was first reported in *Arabidopsis clsy* single mutants along with expected sRNA-dependent hypomethylation (Yang et al. 2018). Presence of hyper-DMRs in *clsy* single mutants indicated that CLSYs regulate global DNA methylation and demethylation balance in the genome. It was postulated that DNA demethylation was possibly defective in *clsy* individual mutants to ensure hypermethylation is maintained. However, expression of ROS1 and other demethylases were unaffected in *Arabidopsis* clsy single mutants. In rice among the six demethylases, *OsROS1* and *OsDML4* were significantly downregulated in rice endosperm in osclsy4-kd. Interestingly, both of the demethylases were documented as crucial players in rice endosperm development (Lang and Gong 2018; Liu et al. 2018; Yan et al. 2022).

In several hyper DMRs, sRNA-independent hypermethylation was observed and the hypermethylated sites were distinct between individual clsy-kd lines. It indicates a possibility that hypermethylation is not necessarily due to redundancy or competition in CLSY-bound regions by individual CLSYs.

Pol IV-suppressed sRNAs that depend on Pol II in rice and *Arabidopsis* led to the production of different-sized sRNAs from diverse genomic regions (Hari Sundar G et al. 2023). Such a regulation required CLSY proteins and Pol IV complex assembly, but not other downstream players including DCLs and AGOs (Hari Sundar G et al. 2023). This suggests that different polymerases compete for specific genomic regions to initiate RdDM through various mechanisms. Blocking the canonical RdDM might promote stronger non-canonical RdDM, resulting in hypermethylation at certain loci (Nuthikattu et al. 2013).

It is also possible that the absence of RdDM in specific regions controlled by individual CLSY proteins might have altered nucleosome positions or chromatin marks to initiate more methylation through their properties as chromatin remodelers. For example, FACT complex protein SSRP1 plays a role in DME-mediated DNA demethylation in *Arabidopsis* endosperm (Ikeda et al. 2011; Frost et al. 2018). Some of the hypermethylation might not be through sRNAs, but through other sRNA-independent DNA methylation or silencing such as mediated in *ddm1* and *mom1* mutants (Vaillant et al. 2006; Saze and Kakutani 2007; Akinmusola et al. 2023). Future research could investigate the mechanism by which hypermethylation arises in these mutants.

It is interesting to note that individual CLSYs compared here target same family of TEs without competing for a single TE in endosperm. Since such a targeting was essential in regulating gene expression necessary for endosperm development and imprinting, it is unclear how such a preference is initiated and maintained. It would be an interesting area for further investigation to understand how TEs and other features are recognized by the CLSYs.

Similar to osclsy3-kd, osclsy4-kd also showed smaller endosperm but in osclsy4-kd, the seeds were slightly wider than PB1. Many known endosperm-development associated genes such as OsBZR1, GW6, OsYUCCA11, NF-YC12, OsDML4, AWN1 and RAG2 were downregulated in osclsy4-kd. Interestingly, osclsy4-kd seeds showed grain chalkiness phenotype which was not observed in osclsy3-kd. This phenotype was also observed in OsCLSY3 OE seeds where OsCLSY3 expression was more than control WT tissues (Pal et al. 2024). This suggests that higher expression of OsCLSY3 beyond a threshold result in chalkiness possibly by altering expression of imprinted, seed development genes or metabolic pathway-related genes. Altered seed size, endosperm quality and developmental timing due to altered dose of expression of imprinted genes is well known among plants. Many well-studied seed size-related and imprinted genes such as Os02g0517531, Os05g0477200, Os05g0447200, Os02g0682200 Os07g0530400, Os05g0490700 were mis-expressed in OsCLSY3 OE, osclsy4-kd and osclsy3-kd endosperm tissues (Supplemental Data 9 and 10). From these observations it appears that proper ratio of OsCLSY3 and OsCLSY4 is absolutely important to maintain proper endosperm development, especially cellularization timing.

The grain chalkiness is a complex phenotype which is a result of water, heat and nutritional stressed conditions in addition to some genetic mutants (Chen et al. 2024). In normal rice grains, starch polygonal granules are tightly arranged, whereas in chalky grains, the granules are spherical and loosely packed. During milling and cooking, chalky grains are easily broken and also affect nutrient value (Chen et al. 2024). Here, we found chalky grains in osclsy4-kd endosperm. Since a direct correlation between the abundance of sRNAs and the expression of shortlisted grain-chalkiness associated genes in osclsy4-kd, multiple severe developmental defects might have collectively contributed to this phenotype. The RdDM pathway helps plants respond to many abiotic stresses such as heat, cold, drought, salt and nutrient starvation (Jiang et al. 2014; Xu et al. 2015; Zhang et al. 2016). TEs are well-known to get de-repressed in abiotic stressed conditions to get inserted near genes. The RdDM is involved in silencing of those TEs to protect genome integrity during stress conditions (Popova et al. 2013). Here, we found grain chalkiness, an environment stress-induced phenotype, is under the regulation of OsCLSY4 mediated RdDM pathway. Future research might reveal how each upstream RdDM pathway players including but not limited to CLSY proteins, regulate gene expression under stresses that crops encounter routinely.

## Methods

### Rice transformation and plant growth

For generating transgenic rice plants, *Agrobacterium*-mediated transformation was performed as described previously (Hiei et al. 1997; Sridevi et al. 2003). Briefly, 3 weeks old scutellum-derived calli from PB1 line (*O. sativa indica*) were used. Calli were infected with *A. tumefaciens* strain LBA4404 having *vir* helper plasmid pSB1 (pSB1 carries extra copies of *vir* genes) and the binary plasmids of interest. Hygromycin antibiotic was used as a selection marker. The regenerated transgenic seedlings were maintained in a growth chamber at 23°C with 16h/8h light/dark cycle at 70 % relative humidity (RH). The plants were subsequently transferred to a greenhouse.

### Vector design and construction

The amiR that targets OsCLSY4 mRNA was designed using the WMD3 web tool (Ossowski et al. 2008; Warthmann et al. 2008), with the stringent criteria for robust amiR generation as described previously (Narjala et al. 2020). The amiR-precursors were synthesized by GeneArt (Thermofisher). The amiR sequence was cloned into pCAMBIA1300 under maize Ubiquitin promoter (P:ZmUbi) with *hygromycin phosphotransferase (hph)* gene as a selection marker. The construct was verified by restriction enzyme-based analysis followed by sequencing. Construct was mobilized into *Agrobacterium* strain LBA4404 (pSB1), and mobilization was verified by PCR analysis.

### Phenotyping of transgenic plants

Phenotypes of transgenic plants such as plant height, panicle length and spikelets number per panicle were measured using (n>6) mature plants grown for about 4 months. Phenotypes of roots and shoots of transgenic plants were measured using (n>8) media-grown 10-day-old seedlings. Images of rice seeds and endosperm from rice lines were obtained using Lecia S8APO stereomicroscope and Nikon camera. For statistical analysis, paired *t*-test was used.

### RNA extraction and RT-qPCR

Total RNA extraction from rice tissues was performed using TRIzol® Reagent (Invitrogen) as per manufacturer’s instructions. For endosperm tissue, RNA isolation was performed as described earlier (Wang et al. 2012; Pal et al. 2024). RT-qPCR was performed for the expression of OsCLSY4 and other genes. First-strand cDNA was synthesized from 1.0 µg of total RNA using Thermo Scientific RevertAid First Strand cDNA Synthesis Kit, as per manufacturer’s instructions. qPCR was carried out with Solis Biodyne - 5x HOT Firepol Evagreen qPCR Master Mix. As internal control, *OsActin* (Os03g0718100) and *Glyceraldehyde-3-phosphate dehydrogenase* (*OsGAPDH)* (Os04g0486600) were used. RT-qPCRs were performed at least three times using the BioRad CFX system. Primers used for analysis are provided in Supplemental Table 3.

### DNA methylation analyses

Total DNA was isolated from 20 day-old endosperm using CTAB method (Rogers and Bendich 1994). Equal amount of DNA was sheared to produce 350 bp fragments using ultrasonication (Covaris). Bisulfite conversion was performed using EZ DNA Methylation-Gold Kit as per the manufacturer’s instructions. The libraries were constructed using the IDT xGen™ Methyl-Seq Lib Prep (Catalog no-10009860) and sequenced using NovaSeq 6000 (2X 100 bp mode). Obtained reads were trimmed using Trimmomatic after quality checking (Bolger et al. 2014). The reads were aligned to the IRGSP1.0 genome using Bismark aligner tool with default parameters (Krueger and Andrews 2011). DNA methylation status was extracted and coverage reports were generated using Bismark tools. The CG, CHG and CHH reports were made from the CX report file. The metaplots and heatmaps were generated using the ViewBS methylation package (Huang et al. 2018).

For the targeted BS sequencing, total DNA was isolated from 60-day-old leaf tissue using the CTAB method. About 200-400 ng of DNA was treated using EZ DNA Methylation-Gold Kit (Zymo Research). The treated DNA was used for PCR as a template. Targeted regions were amplified by JumpStart™ Taq DNA Polymerase (Sigma). The PCR products were deep sequenced on NovaSeq 6000 (2X 100 bp mode) platform. The obtained reads were quality checked and trimmed using cutadapt (Martin 2011) and aligned to create genome (target sites) using Bismark aligner tool (Krueger and Andrews 2011). The obtained results are analyzed using methylation package ViewBS (Huang et al. 2018). Primers used for analysis are provided in Supplemental Table 3.

### sRNA library preparation and differential expression analyses

Total RNA extraction from endosperm and sRNA library preparations were carried out as described previously (Hari Sundar G et al. 2023). After checking the quality of obtained reads, the adaptors were trimmed by UEA sRNA Workbench (Stocks et al. 2018). The filtered sRNAs were classified into 21-22 nt and 23-24 nt reads, and aligned using Bowtie -v 1 -m 100 -y -a --best –strata (Langmead et al. 2009). The sRNAs loci were identified using Shortstack with the following parameters: --nohp --mmap f -- mismatches 1 -mincov 5 rpm (Axtell 2013). CLSY-dependent sRNA loci were identified by bedtools intersect (Quinlan and Hall 2010). For quantifying the sRNA abundance from transposons, and siren loci, bedtools multicov was used to obtain raw abundance and then normalized to RPKM values (Quinlan and Hall 2010; Quinlan 2014). Obtained values were plotted as box plots using custom R scripts in ggplot2 (Wickham 2016). The venn diagrams were generated by intervene online tools (Khan and Mathelier 2017). For publicly available datasets sRNAs loci were identified using Shortstack with following parameters: --nohp --mmap f --mismatches 1 -mincov 2 rpm (Axtell 2013).

### RNA-seq and analyses

RNA-seq was performed using 20-day-old endosperm tissues as source tissue. Poly(A) enrichment was performed before the library preparation using NEBNext® Ultra™ II Directional RNA Library Prep kit (E7765L) as per manufacturer’s instructions. The obtained libraries were sequenced in NovaSeq 6000 (2X 100 bp mode) platform. The obtained reads were adapter trimmed using Trimmomatic after the quality checking (Bolger et al. 2014) and aligned to IRGSP1.0 genome using HISAT2 with default parameters (Kim et al. 2015). Cufflinks was used to perform differential gene expression (DEG) analyses and statistical testing (Trapnell et al. 2012). Volcano plots were generated for DEGs using custom R scripts with ap-value cut-off of less than 0.05 and the absolute log_2_ (fold change) expression cut-off of more than 1.5. For quantifying the expression of genes, bedtools multicov (Quinlan and Hall 2010; Quinlan 2014; Patwardhan et al. 2019) was used to obtain raw abundance and then normalized to RPKM values. DEseq2 was used to identify log_2_1.5 fold change upregulated and downregulated genes with the p-value cut-off of less than 0.05 (Love et al. 2014). These values were plotted as box plots and heatmaps using custom R scripts in ggplot2 (Wickham 2016). For publicly available datasets, log_2_1.0 fold change upregulated and downregulated genes with the p-value cut-off of less than 0.05 were identified by DEseq2 (Love et al. 2014).

### GO analysis

The GO analysis was performed using ShinyGO v0.75 platform (Ge et al. 2020). The gene IDs were used from RAPDB. The biological processes with FDR cutoff of p-value-0.05.

### SEM imaging

For scanning electron microscope (SEM) imaging, rice endosperms were collected (25 DAP) and fixed in 16 % Formaldehyde, 25 % Glutaraldehyde and 0.2 M cacodylate buffer for 12-16 h. The samples were rinsed with double distilled water and dehydrated in a series of ethanol (30%,50%,70% and 100%) passages and dried in critical point drying (CPD, Leica EM CPD300). After CPD, endosperms were cut by a sharp razor blade. Samples were coated with gold, and the images were obtained using a Carl Zeiss scanning EM (Scanner SE2) at an accelerating voltage of 2 kV-4 kV as described before (Pachamuthu et al. 2021; Hari Sundar G et al. 2023).

### Germination assay

The germination assay was performed as described previously (He et al. 2020). A total of 20 seeds of different genotypes (n=5) were imbibed on the wet filter paper in 5 cm diameter petri-plates. In all the plates, 4 ml of single distilled water was used to wet the filter papers and seeds were germinated at 25°C for 7 days in the dark.

Experiments were repeated twice independently. The germination assay was also performed in MS media with surface sterilized seeds of different genotypes.

### Pollen staining

Pollen viability test was performed using I_2_-KI staining solution containing 0.2% (w/v) I_2_ and 2% (w/v) KI as described (Khatun and Flowers 1995; Yao et al. 2018; Pachamuthu et al. 2021). Anthers from six spikelets of mature panicles were collected in 200 µl of solution one day before the fertilization. Mechanical shearing was used to release pollens into the solutions. After 10 min, viable pollen grains were counted under the bright-field microscope (OLYMPUS BX43). Before imaging, anther and other debris were removed carefully. Dark stained and round pollen grains were considered as viable, while very light blue and distorted pollens were considered as non-viable (Das et al. 2020; Hari Sundar G et al. 2023).

### Statistics and reproducibility

No data were excluded from the analysis. All SEM images were performed at least twice with different samples. Statistical analysis was performed using two-tailed paired Student’s *t-*test or two-sided Wilcoxon test to determine differences between two groups. Statistical analyses were performed using excel and R studio. Details such as the number of replicates, the level of significance and sample sizes for RT-qPCR, RNA analysis, sequencing, phenotyping were mentioned in the corresponding figure legends, text and Supplemental Data. A full list of primers, probes, sequences and other details are available in Supplemental Tables.

## Data availability

All raw and processed sequencing data generated in this study have been submitted to the NCBI Gene Expression Omnibus (GEO; https://www.ncbi.nlm.nih.gov/geo/) under accession numbers GSEXXXXXX.

## Acknowledgements

We thank lab members for comments and suggestions. We thank Prof. K. Veluthambi for *Agrobacterium* strains, PB1 seeds and binary plasmids. We thank genomics, electron microscopy, IT, radiation, greenhouse, and lab kitchen facilities at the NCBS. This work was supported by NCBS-TIFR core funding, Government of India. This study was also supported by the Department of Atomic Energy, Government of India, under Project Identification No. RTI 4006 (1303/3/2019/R&D-II/DAE/4749 dated 16.7.2020). These funding agencies did not participate in the design of experiments, analysis, or interpretation of data or in writing the manuscript. We thank anonymous reviewers for their valuable suggestions and insightful comments.

## Author contributions

PVS and AKP designed all experiments and discussed results and wrote the manuscript. AKP performed most of the experiments and bioinformatics analysis. SR and RD performed phenotype and transgene analysis. All authors have read and approved the manuscript.

