## Supplementary Figures and Tables for "Loss of function of chromatin remodeler OsCLSY4 leads to RdDM-mediated mis-expression of endosperm-specific genes affecting grain qualities"

**Supplemental Figures 1 to 7**

**Supplemental Tables 1 to 3**

### Supplemental information

This manuscript has 7 supplemental figures, 3 supplemental tables and 10 Supplemental Datasets.

### Supplemental Figures

**Supplemental Figure S1.** Phenotypes of osclsy4-kd plants.

**Supplemental Figure S2.** Endosperm-sRNAs are regulated by both OsCLSYs.

**Supplemental Figure S3.** CLSYs regulates expression of 21-22 nt sRNAs in endosperm.

**Supplemental Figure S4.** DNA methylation of an RdDM loci in leaf, regulated by OsCLSY4.

**Supplemental Figure S5.** CLSYs regulate DNA methylation in endosperm non-redundantly.

**Supplemental Figure S6.** OsCLSY4 regulates expression of protein coding genes.

**Supplemental Figure S7.** OsCLSY4 target different regions for sRNA production in seedling.

### Supplemental Tables

**Supplemental Table 1:** Details of high-throughput genomics data generated in this study.

**Supplemental Table 2:** Details of high-throughput genomics data obtained from publicly available datasets.

**Supplemental Table 3:** List of oligos and probes used in this study.

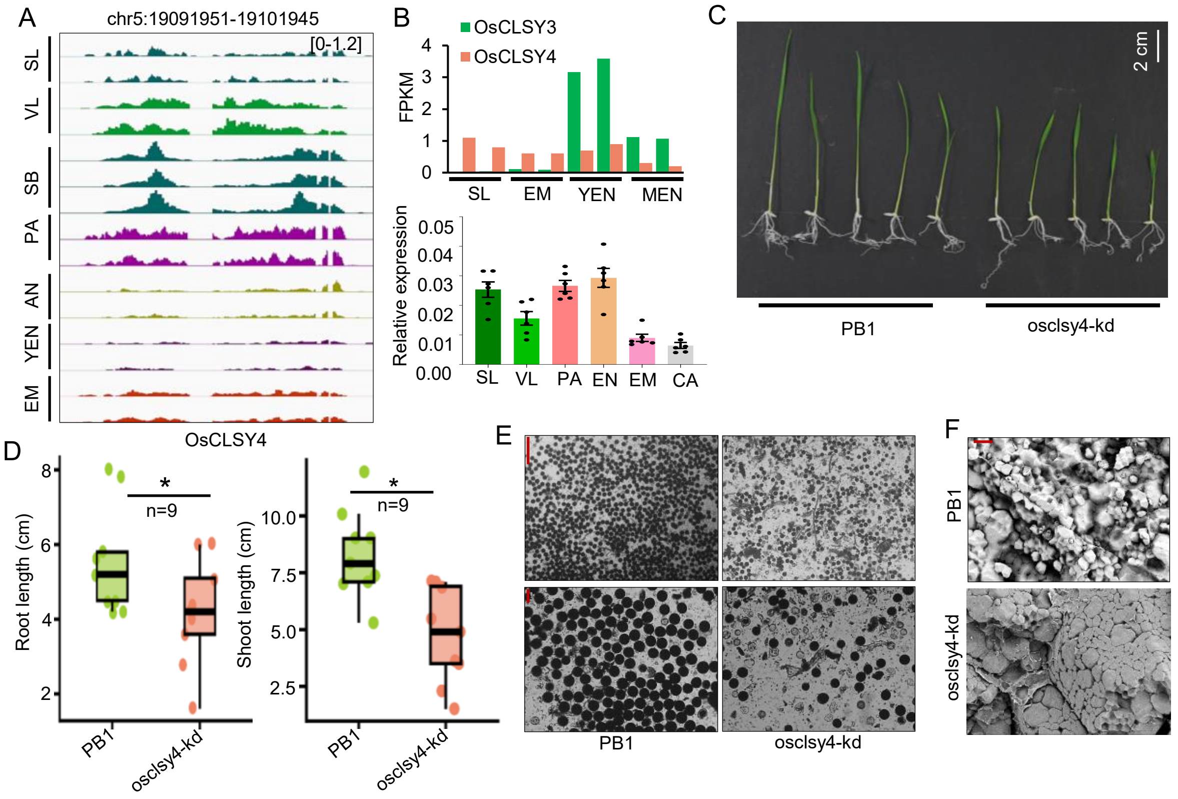


**Supplemental Figure S1. Phenotypes of osclsy4-kd plants.**
(A) IGV screenshot representing expression of OSCLSY4 gene across different rice tissues. SL-seedling (GSE130168), VL- vegetative leaf (GSE138705), SB-shootbase (GSE131319), PA-panicle (GSE180457), AN-anther (GSE180457), YEN-young endosperm (GSE229959), EM-embryo (GSE229959). (B) Barplots showing expression of OsCLSY4 and OsCLSY3 across different rice tissues. MEN-Mature endosperm (GSE229959) in transcriptomes (top). RTqPCR analysis showing expression of OsCLSY4 in different tissues, CA-calli. *OsGAPDH* served internal control. (C) Image showing phenotypes of PB1 and osclsy4-kd 10 days old seedlings. (D) Boxplots showing root and shoot length of PB1 and osclsy4-kd seedlings. (E) Pollen viability assay in PB1 and osclsy4-kd. Scale bar (SB)-100 µm. (F) Electron microscopy images of PB1 and osclsy4-kd EN. SB-10 µm.

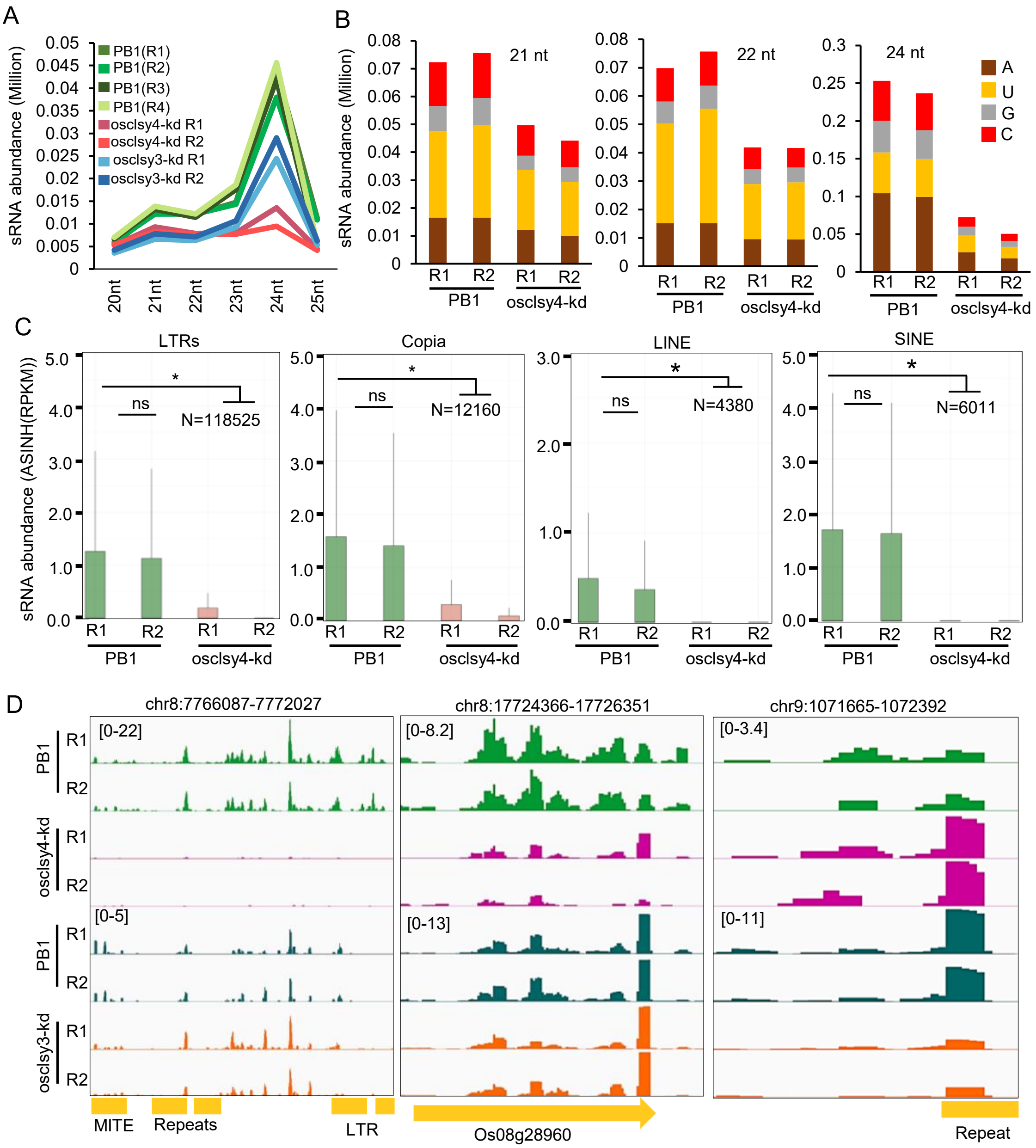


**Supplemental Figure S2. Endosperm-sRNAs are regulated by both OsCLSYs.**
(A) Plot showing expression of 21-25 nt sRNAs in PB1 and individual kd lines in EN. (B) Stacked barplot showing abundance of first 5' nucleotide of mapped sRNAs in WT and clsy4-kd EN. (C) Boxplots representing expression of 23-24 nt sRNAs across different TEs. *-significant. ns-non-significant (Wilcoxon test $p < 0.01$). (D) IGV screenshots showing expression of 23-24 nt sRNAs in different CLSY-dependent sRNA loci.

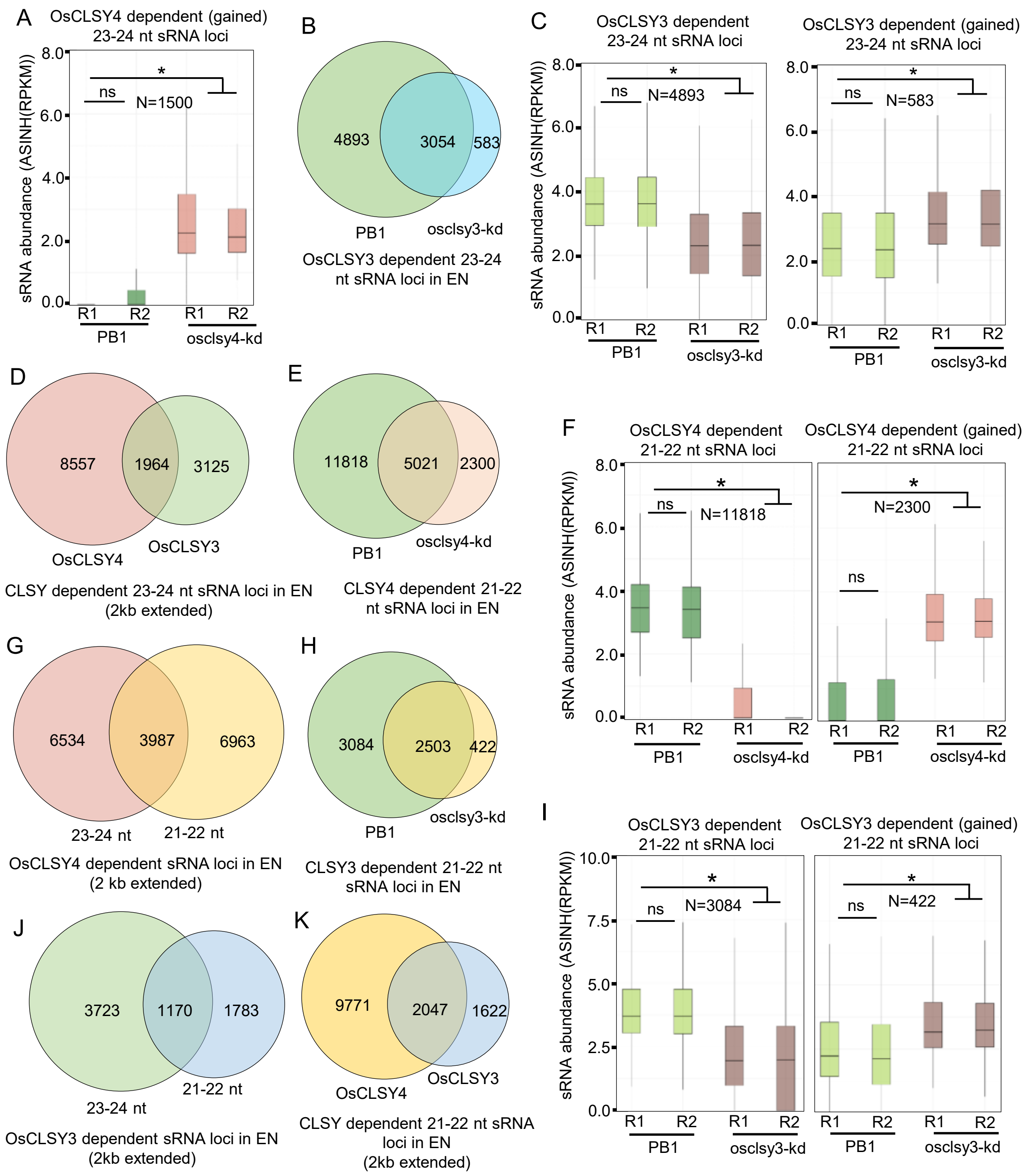


**Supplemental Figure S3. CLSYs regulates expression of 21-22 nt sRNAs in endosperm.** (A) Boxplots representing OsCLSY4 dependent 23-24 nt gained sRNA loci in EN. (B) Venn diagram showing OsCLSY3 dependent 23-24 nt sRNA loci in EN. (C) Boxplots representing abundance of 23-24 nt sRNAs from OsCLSY3 dependent 23-24 nt sRNA loci (lost and gained, respectively). (D) Venn diagram showing overlap between OsCLSY4 and OsCLSY3 dependent 23-24 nt sRNA loci (1 kb extended both sides). (E) Venn diagrams representing OsCLSY4-dependent 21-22 nt sRNAs. (F) Boxplots representing abundance of 21-22 nt sRNAs from OsCLSY4 dependent 21-22 nt sRNA loci (lost and gained, respectively). *-significant. ns-non-significant (Wilcoxon test $p < 0.01$). (G) Venn diagram showing overlap between OsCLSY4 dependent 21-22 nt and 23-24 nt sRNA loci (1 kb extended on both sides). (H) Venn diagrams representing OsCLSY3-dependent 21-22 nt sRNAs. (I) Boxplots representing abundance of 21-22 nt sRNAs from OsCLSY3 dependent 21-22 nt sRNA loci (lost and gained, respectively). (J) and (K) Venn diagrams representing overlap between OsCLSY4 dependent 21-22 nt, 23-24 nt sRNA loci (1 kb extended both sides) and overlap between OsCLSY3 and OsCLSY4 dependent 21-22 nt sRNA loci, respectively (1 kb extended on both sides).

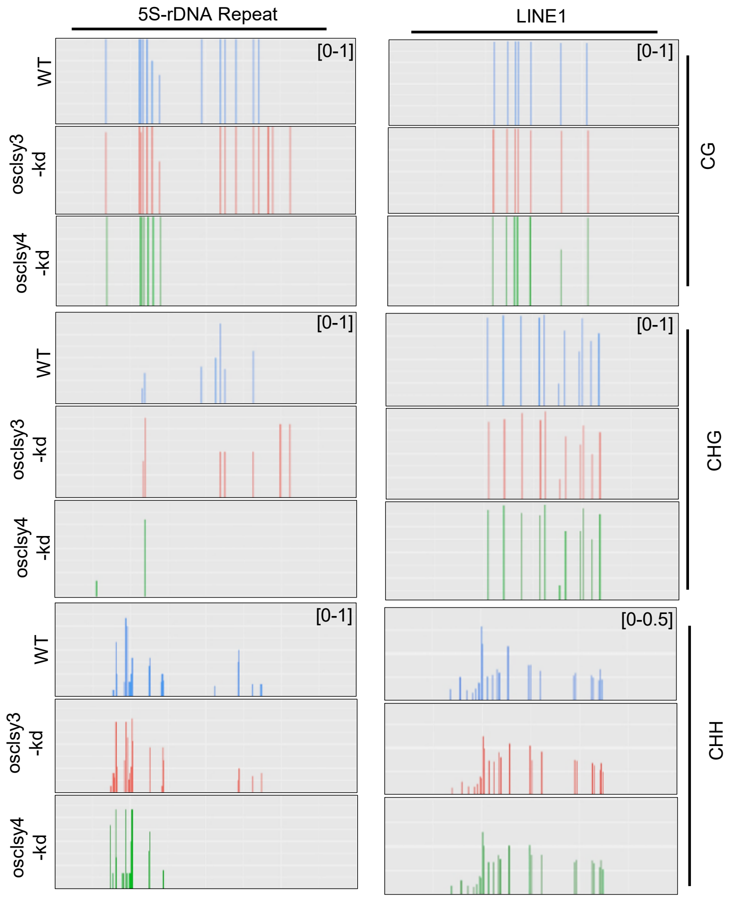


**Supplemental Figure S4. DNA methylation of an RdDM loci in leaf, regulated by OsCLSY4.**
Targeted bisulfite PCR showing DNA methylation of 5S rDNA repeats and LINE1 TEs in osclsy3-kd and osclsy4-kd leaves.

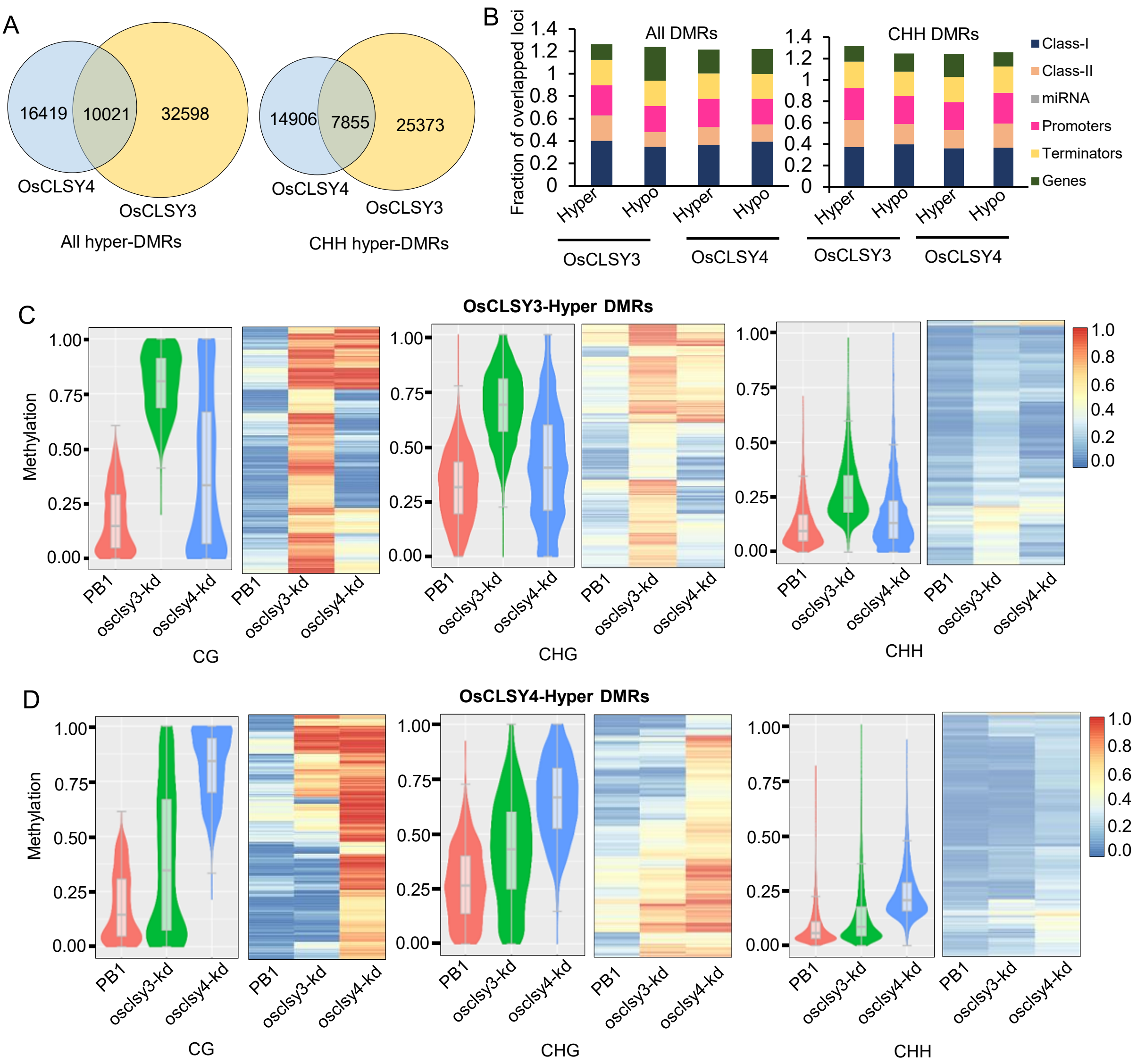


**Supplemental Figure S5. CLSYs regulate DNA methylation in endosperm non-redundantly.** (A) Venn diagrams representing overlap between OsCLSY4 and OsCLSY3 hyper-DMRs. (B) Plots depicting genomic features of OsCLSYs hypo- and hyper-DMRs. (C), (D) Violin-plots and heatmaps showing OsCLSY3 and OsCLSY4-dependent hyper-DMRs, respectively.

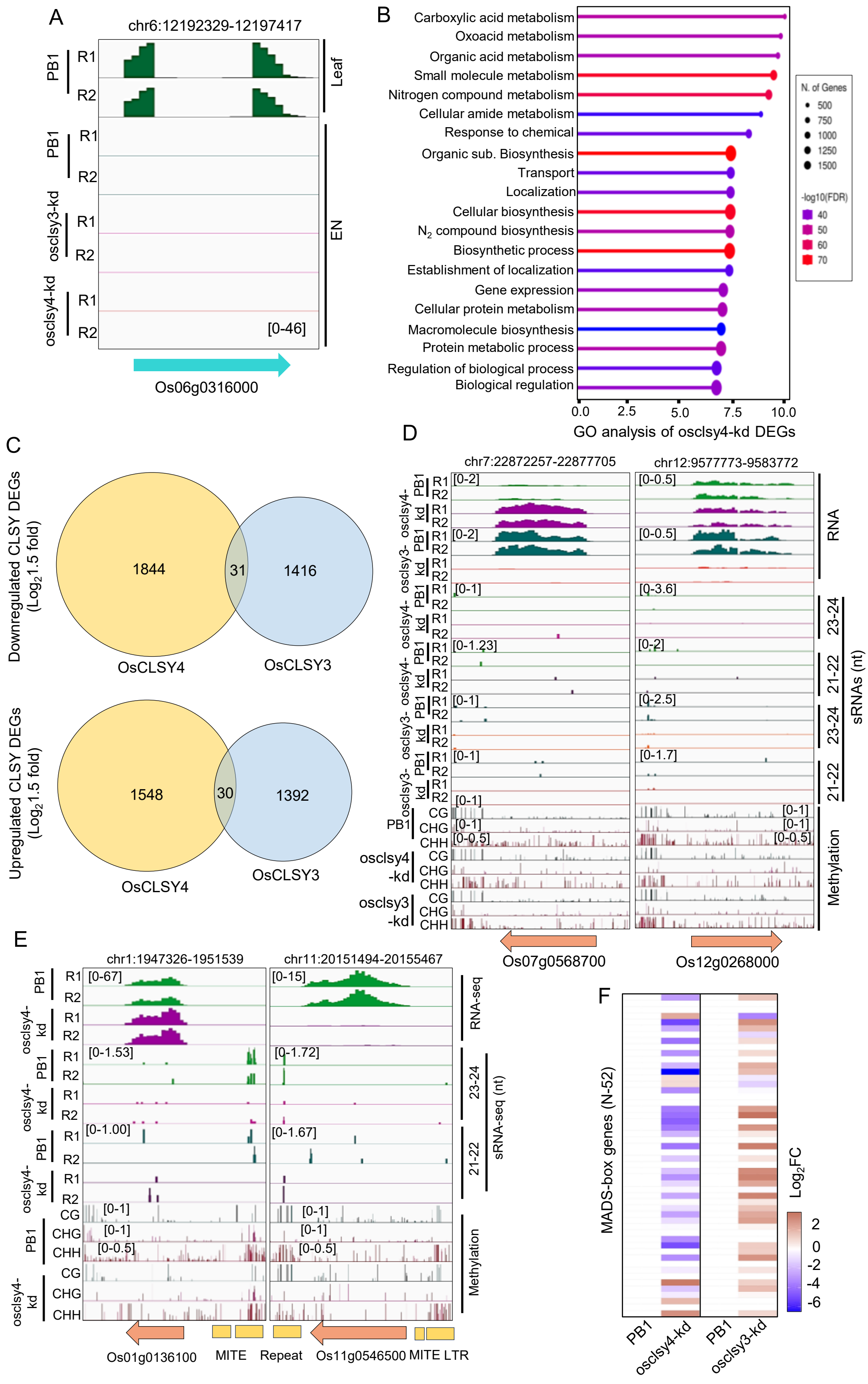


**Supplemental Figure S6. OsCLSY4 regulates expression of protein coding genes.** (A) IGV screenshots showing lack of expression of green tissue-specific gene in EN transcriptomes. (B) GO analysis of osclsy4-kd EN. (C) Venn diagrams showing overlap between downregulated and upregulated DEGs in genotypes. (D) IGV screenshots showing expression of two DEGs in osclsy3-kd and osclsy4-kd EN. (E) IGV screenshots showing expression of an upregulated and downregulated genes in clsy4-kd EN. (F) Heatmap showing expression of MADS-box genes in osclsy4-kd and osclsy3-kd EN.

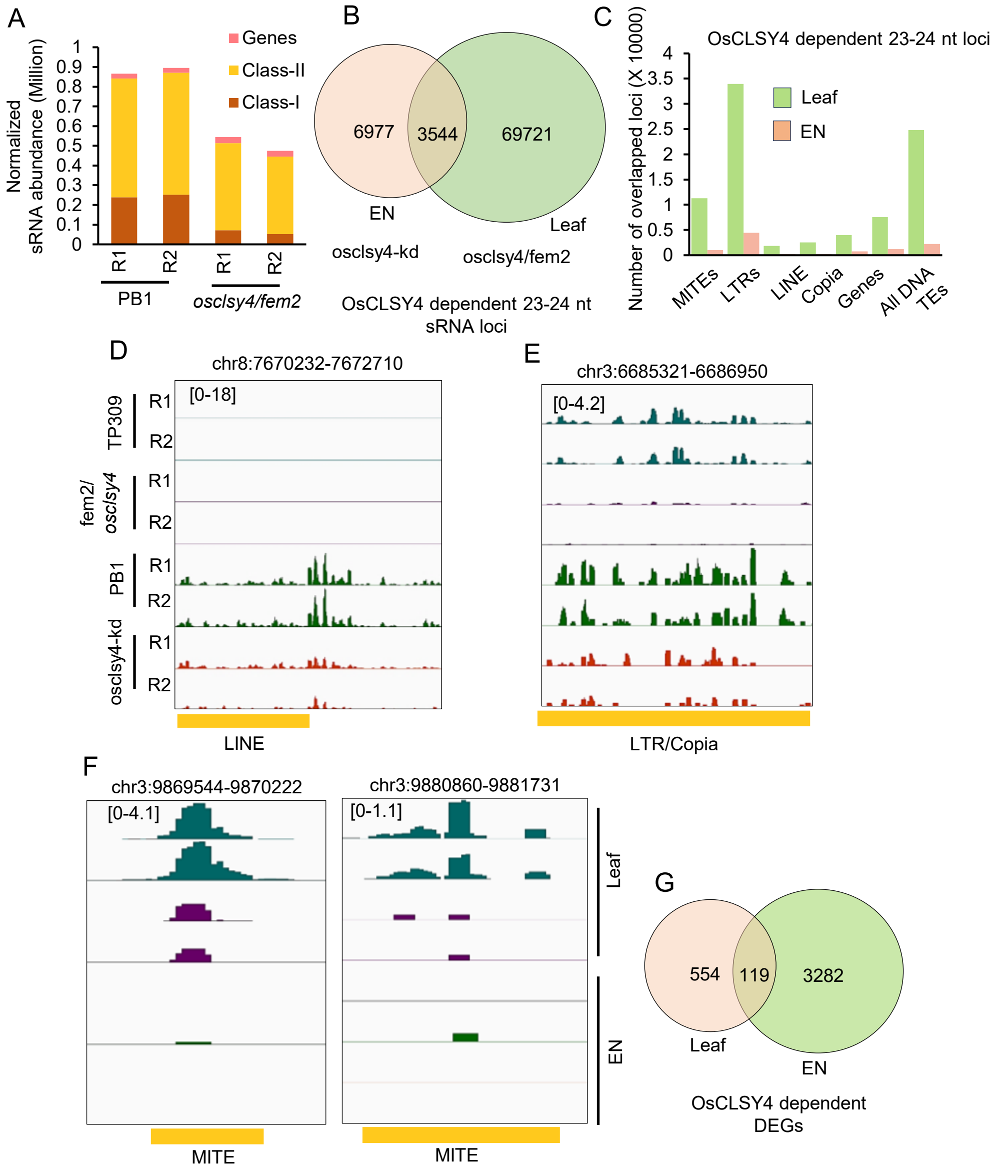


**Supplemental Figure S7. OsCLSY4 target different regions for sRNA production in seedling. and endosperm tissues.** (A) Plot showing 23-24 nt sRNA abundance across different genomic features in osclsy4/fem2 seedling tissue. (B) Venn diagrams showing overlap between OsCLSY4 dependent 23-24 nt sRNA loci in EN and leaf. (C) Bar plot representing the overlap of OsCLSY4-dependent 23-24 nt sRNA loci with distinct genomic regions. (D),(E), (F) IGV screenshots showing sRNA expression in TEs and repeats in leaf and EN in fem2 and osclsy4-kd lines. (G) Venn diagram showing overlap between fem2 and osclsy4-kd DEGs.

Supplementary Table 1: Details of high-throughput genomics data generated in this study

| Sl. No | Dataset type | Genotype | Replicate | Source tissue | GSM Number | GSE Number | Total number of mapped reads obtained | Sequencing mode |
|---|---|---|---|---|---|---|---|---|
| 1 | Small RNA-seq | WT (PB1) | Rep1 | 20 days Endosperm | GSM8790179 | GSE289382 | 19374449 | Paired end |
| 2 | Small RNA-seq | WT (PB1) | Rep2 | 20 days Endosperm | GSM8790180 | GSE289382 | 16241812 | Paired end |
| 3 | Small RNA-seq | osclsy4-kd | Rep1 | 20 days Endosperm | GSM8790181 | GSE289382 | 21233703 | Paired end |
| 4 | Small RNA-seq | osclsy4-kd | Rep2 | 20 days Endosperm | GSM8790182 | GSE289382 | 19169541 | Paired end |
| 5 | RNA-seq | WT (PB1) | Rep1 | 20 days Endosperm | GSM8785346 | GSE289150 | 26610703 | Paired end |
| 6 | RNA-seq | WT (PB1) | Rep2 | 20 days Endosperm | GSM8785347 | GSE289150 | 26447919 | Paired end |
| 7 | RNA-seq | osclsy4-kd | Rep1 | 20 days Endosperm | GSM8785348 | GSE289150 | 24060165 | Paired end |
| 8 | RNA-seq | osclsy4-kd | Rep2 | 20 days Endosperm | GSM8785349 | GSE289150 | 27848043 | Paired end |
| 9 | RNA-seq | OsCLSY3OE | Rep1 | 20 days Endosperm | GSM8785350 | GSE289150 | 12797491 | Paired end |
| 10 | RNA-seq | OsCLSY3OE | Rep2 | 20 days Endosperm | GSM8785351 | GSE289150 | 11505785 | Paired end |
| 11 | Targeted bisulfite | WT Leaf (PB1) | Rep1 | 60 days Leaf | GSM8785386 | GSE289154 | 528089 | Paired end |
| | Targeted bisulfite | WT Leaf (PB1) | Rep2 | 60 days Leaf | GSM8785387 | GSE289154 | 570046 | Paired end |
| 12 | Targeted bisulfite | osclsy4-kd Leaf | Rep1 | 60 days Leaf | GSM8785388 | GSE289154 | 698626 | Paired end |
| | Targeted bisulfite | osclsy4-kd Leaf | Rep2 | 60 days Leaf | GSM8785389 | GSE289154 | 549683 | Paired end |
| 13 | Targeted bisulfite | osclsy3-kd Leaf | Rep1 | 60 days Leaf | GSM8785390 | GSE289154 | 567914 | Paired end |
| | Targeted bisulfite | osclsy3-kd Leaf | Rep2 | 60 days Leaf | GSM8785391 | GSE289154 | 513113 | Paired end |
| 14 | Bisulfite-Seq | osclsy4-kd Endosperm | NA | 20 days Endosperm | GSM8785379 | GSE289152 | 87951323 | Paired end |

Supplementary Table 2: Details of high-throughput genomics data obtained from publicly available datasets

| Sl. No | Dataset type | Genotype | Source tissue | SRA number | GSE number | Reference |
|---|---|---|---|---|---|---|
| 1 | RNAseq | WT (PB1) | pre-emerged panicle | SRX11493038 | GSE180457 | [1] |
| 2 | RNAseq | WT (PB1) | pre-emerged panicle | SRX11493039 | GSE180457 | [1] |
| 3 | RNAseq | WT (PB1) | Anther | SRX11493042 | GSE180457 | [1] |
| 4 | RNAseq | WT (PB1) | Anther | SRX11493043 | GSE180457 | [1] |
| 5 | RNAseq | WT_T1_r1 | Leaf | SRX6976682 | GSE138705 | [2] |
| 6 | RNAseq | WT_T1_r2 | Leaf | SRX6976683 | GSE138705 | [2] |
| 7 | RNAseq | Nip_se_rep1 | Seedling | SRX5724238 | GSE130168 | [3] |
| 8 | RNAseq | Nip_se_rep2 | Seedling | SRX5724239 | GSE130168 | [3] |
| 9 | RNAseq | WT_base_rep1 | Shoot base of seedling | SRX5846194 | GSE131319 | [4] |
| 10 | RNAseq | WT_base_rep2 | Shoot base of seedling | SRX5846195 | GSE131319 | [4] |
| 11 | RNAseq | WT_base_rep3 | Shoot base of seedling | SRX5846196 | GSE131319 | [4] |
| 12 | RNA-seq | Embryo_Rep1 | 25 days Embryo | SRX20001598 | GSE229959 | [5] |
| 13 | RNA-seq | Embryo_Rep2 | 25 days Embryo | SRX20001599 | GSE229959 | [5] |
| 14 | RNA-seq | Mature_endosperm_Rep1 | 25 days Endosperm | SRX20001600 | GSE229959 | [5] |
| 15 | RNA-seq | Mature_endeosperm_Rep2 | 25 days Endosperm | SRX20001601 | GSE229959 | [5] |
| 16 | RNA-seq | Young_endosperm_Rep1 | 15 days Endosperm | SRX20001602 | GSE229959 | [5] |
| 17 | RNA-seq | Young_endosperm_Rep2 | 15 days Endosperm | SRX20001603 | GSE229959 | [5] |
| 18 | RNA-seq | WT_CLSY3_Endosperm_Rep1 | 20 days Endosperm | SRX20001606 | GSE229959 | [5] |
| 19 | RNA-seq | WT_CLSY3_Endosperm_Rep2 | 20 days Endosperm | SRX20001607 | GSE229959 | [5] |
| 20 | RNA-seq | KD_CLSY3_Endosperm_Rep1 | 20 days Endosperm | SRX20001604 | GSE229959 | [5] |

| | | | | | | |
|---|---|---|---|---|---|---|
| 21 | RNA-seq | KD_CLSY3_Endosperm_Rep2 | 20 days Endosperm | SRX20001605 | GSE229959 | [5] |
| 22 | RNA-seq | Nip, rep3 | 18 days old seedling | SRX17907626 | GSE215853 | [6] |
| 23 | RNA-seq | Nip, rep4 | 18 days old seedling | SRX17907627 | GSE215853 | [6] |
| 24 | RNA-seq | fem2-3, rep1 | 18 days old seedling | SRX17907628 | GSE215853 | [6] |
| 25 | RNA-seq | fem2-3, rep2 | 18 days old seedling | SRX17907629 | GSE215853 | [6] |
| 26 | RNA-seq | fem2-3, rep3 | 18 days old seedling | SRX17907630 | GSE215853 | [6] |
| 27 | Small RNA-seq | WT_CLSY3_Endosperm_Rep1 | 20 days Endosperm | SRX20001558 | GSE229958 | [5] |
| 28 | Small RNA-seq | WT_CLSY3_Endosperm_Rep2 | 20 days Endosperm | SRX20001559 | GSE229958 | [5] |
| 29 | Small RNA-seq | KD_CLSY3_Endosperm_Rep1 | 20 days Endosperm | SRX20001556 | GSE229958 | [5] |
| 30 | Small RNA-seq | KD_CLSY3_Endosperm_Rep2 | 20 days Endosperm | SRX20001557 | GSE229958 | [5] |
| 31 | Small RNA-seq | TP309_seedling_R1 | 18 days old seedling | SRX11502899 | GSE130166 | [3] |
| 32 | Small RNA-seq | TP309_seedling_R2 | 18 days old seedling | SRX11502900 | GSE130166 | [3] |
| 33 | Small RNA-seq | fem2-1 | 18 days old seedling | SRX17907624 | GSE215854 | [3] |
| 34 | Small RNA-seq | fem2-3 | 18 days old seedling | SRX17907625 | GSE215854 | [3] |
| 35 | Bisulfite-Seq | WT_Endosperm | 20 days Endosperm | SRX23802523 | GSE260651 | [5] |
| 36 | Bisulfite-Seq | clsy3-kd_Endosperm | 20 days Endosperm | SRX23802524 | GSE260651 | [5] |
| 37 | Bisulfite-Seq | TP309_seedling_BS | 18 days old seedling | SRX9211924 | GSE158710 | [7] |
| 38 | Bisulfite-Seq | GAS_seedling_BS | 18 days old seedling | SRX9211925 | GSE158710 | [7] |
| 39 | Bisulfite-Seq | fem2-1_BS-seq | 18 days old seedling | SRX17907644 | GSE215855 | [3] |
| 40 | Bisulfite-Seq | fem2-3_BS-seq | 18 days old seedling | SRX17907645 | GSE215855 | [3] |

Supplementary Table 3: List of oligos and probes used in this study

| Oligo Name | Oligo ID | Oligo sequence (5'-3') | Purpose | Reference |
|---|---|---|---|---|
| STN_OsRAC1_F | 2563 | GCTATGTACGTCGCCATCCAGG | RT-qPCR | [1] |
| STN_OsRAC1_R | 2564 | TGAGATCACGCCCAGCAAGG | RT-qPCR | [1] |
| STN_OsGAPDH_qPCR_F | 3873 | GGGTATTCTGGGTTACGTTGAGGAG | RT-qPCR | [1] |
| STN_OsGAPDH_qPCR_R | 3874 | ACGGATCAGGTCAACAACGCGAGAG | RT-qPCR | [1] |
| AP_CLSY4_RT_F | 3419 | gtggtggatccagtgaatgaagagttgg | RT-qPCR | [1] |
| AP_CLSY4_RT_R | 3420 | cccaaaattatcaccgtcgtcatcagc | RT-qPCR | [1] |
| miRNA168 | 32 | gtcgccgagaagatcctccatc | sRNA northern | [2] |
| U6_probes | 13 and 14 | ggccatgctaatcttctctgtatcgtt and ccaattttatcggatgtccccgaagggac | sRNA northern | [2] |
| MITE siRNA | 3430 | ggtcccacctgtcatacacacact | sRNA northern | [2] |
| AP_CLSY4_amiR1_probe | 3880 | GAGAGGTCATTCGTGCTTACA | sRNA northern | [1] |
